# Ion Permeation, Selectivity, and Electronic Polarization in Fluoride Channels

**DOI:** 10.1101/2021.12.08.471811

**Authors:** Zhi Yue, Zhi Wang, Gregory A. Voth

## Abstract

Fluoride channels (Fluc) export toxic F^−^ from the cytoplasm. Crystallography and mutagenesis have identified several conserved residues crucial for fluoride transport, but the permeation mechanism at the molecular level has remained elusive. Herein we have applied constant-pH molecular dynamics and free energy sampling methods to investigate fluoride permeation through a Fluc protein from *Escherichia coli*. We find that fluoride is facile to permeate in its charged form, i.e., F^−^, by traversing through a non-bonded network. The extraordinary F^−^ selectivity is gained by the hydrogen-bonding capability of the central binding site and the Coulombic filter at the channel entrance. The F^−^ permeation rate calculated using an electronically polarizable force field is significantly more accurate compared to the experimental value than that calculated using a more standard additive force field, suggesting an essential role for electronic polarization in the F^−^ – Fluc interactions.

**SIGNIFICANCE**
A comprehensive atomistic-level computational study is presented of the mechanism of fluoride permeation through fluoride channels. The mechanism of fluoride permeation of the F^−^ anion is established and the microscopic determinants of F^−^ selectivity revealed. The essential nature of electronic polarization during F^−^ permeation is also demonstrated through the computational modeling.

## INTRODUCTION

Fluoride is ubiquitous in the biosphere at 20 – 100 µM base levels (1). To alleviate fluoride toxicity (2,3), many organisms utilize transmembrane proteins to reduce cytoplasmic F^−^ accumulation: F^−^/H^+^ antiporters (CLC^F^) found in eubacteria (4), and more broadly distributed F^−^ channels (Fluc, also known as CrcB in bacteria or FEX in eukaryotes) (5).

The Fluc family’s function (6-14) and structure (10,11,15-19) have been studied extensively since its discovery (20-24). Though expressed as a homodimer, two bacterial Fluc monomers work independently so that inactivating one does not block F^−^ conduction through the other (10). Fluc adopts a “dual topology” architecture, which means one monomer inserts into the membrane with its termini facing in and the other monomer inserts into the membrane with its termini facing out (**Fig. 1A**) (6,15,16,21). But the monomers with antiparallel orientation transfer F^−^ in the same direction, as the single-channel conductance is about half of the double-channel value (10). Fluc has a measured turnover rate of 10^6^ – 10^7^ ions/s (6,8), a rate typical for ion channels. Notably, Fluc is exceptionally selective towards F^−^ over Cl^−^ and small cations (6,8).

**FIGURE 1.**
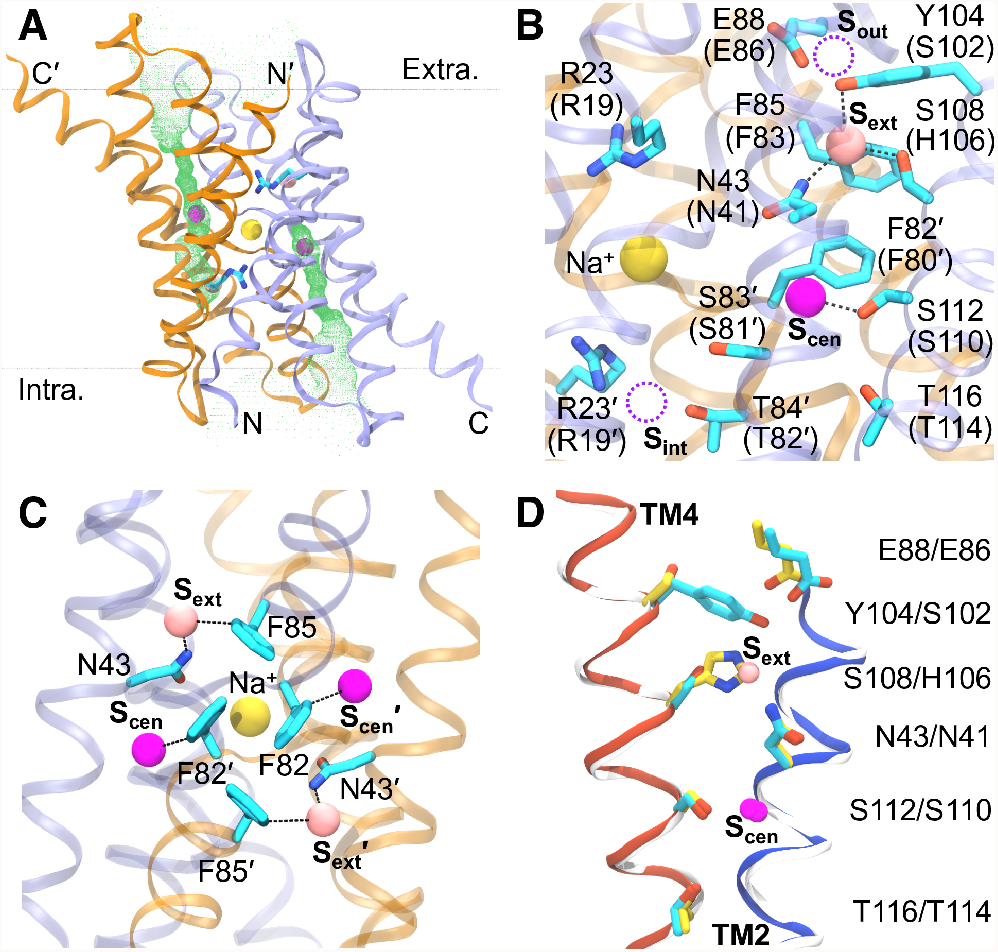
Fluc structure and essential residues. (**A**) Fluc architecture. Fluc-Bpe dimer is shown as ribbon (ice-blue and orange for chains A and B, respectively). Ions are represented by spheres (Na^+^: yellow; S_cen_ F^−^: magenta; S_out_ F^−^: pink). R23s are displayed as sticks. Green mesh depicts the pores generated by HOLE (25). Black dots indicate the lipid head groups calculated by the PPM webserver (26). Bilayer orientation and Fluc termini are labeled. Primed labels belong to chain B. (**B**) F^−^-binding sites. Essential residues and ions are labeled. Parenthesized are residues in Fluc-Ec2. Black dashed lines indicate HBonds. Note S_int_ and S_out_ (dashed purple circles) were seen in Fluc-Ec2. **(C)** “Phe-box”. Black dashed lines indicate HBonds (with N43s) or edgewise anion-aromatic interactions (with F82s/F85s). (**D**) “Polar track”. Helices 2 and 4 from Fluc-Bpe (blue and red) and Fluc-Ec2 (white) are overlaid. Relevant residues (cyan and yellow for Fluc-Bpe and Fluc-Ec2 carbon atoms, respectively) and F^−^ ions are labeled. Note the S_ext_ F^−^ is seen in Fluc-Bpe. The presentation styles adopted here are used throughout the paper unless otherwise stated.

Crystal structures from *Escherichia coli*. (Fluc-Ec2, PDB: 5A43) and *Bordetella pertussis* (Fluc-Bpe, PDB: 5NKQ) have revealed an antiparallel dimer with two segregated F^−^ pores (**Fig. 1A**) (16). In the 5NKQ structure, each monomer harbors two F^−^ ions. One F^−^ is seen in the center (hence denoted as S_cen_) interacting with a conserved F82 (F80 in Fluc-Ec2) from its twin monomer via edgewise anion-aromatic interaction (**Fig. 1B**). The second F^−^ is situated near the pore exit (hereafter termed as S_ext_) and stabilized via hydrogen-bond (HBond) with a strongly conserved N43 (N41 in Fluc-Ec2) and anion-aromatic interaction with a conserved F85 (F83 in Fluc-Ec2). The 5A43 structure only has two S_cen_ F^−^ ions. The side-chains of the four phenylalanines side-to-face assemble into a symmetrical motif (“Phe-box”, **Fig. 1C**) that has been demonstrated to be essential for fluoride permeation (10,16). Beyond S_ext_, there is another binding site S_out_ in some Fluc-Ec2 structures, where F^−^ sits near E86 (E88 in Fluc-Bpe, **Fig. 1B**). The fourth binding site S_int_ at the pore entrance is observed in Fluc-Ec2 mutants crystallized with Br^-^ (19). S_int_ anions interact with twin monomer via HBonds with the side-chain hydroxyls of an invariant S81 (S83 in Fluc-Bpe) and a highly conserved T82 (T84 in Fluc-Bpe), as well as ion-pair (or salt-bridge) with a universally conserved R19 (R23 in Fluc-Bpe) (**Fig. 1B**). **Table S1** of the Supporting Material summarizes the Fluc crystal structures obtained to date. Modestly conserved polar residues on the fourth helix (“polar track”, Y104/S108/S112/T116 in Fluc-Bpe, S102/H106/S110/T114 in Fluc-Ec2, **Fig. 1D**) are expected to further stabilize F^−^ along the pore. In addition, a Na^+^ cation was found deeply buried at the dimer interface and sandwiched by R23s (**Fig. 1AB**). This important Na^+^ (14) was thought to stabilize the dimer and add electropositivity to the pores (5,16).

Nevertheless, the fluoride permeation mechanism remains elusive. Given the HBond donors lining the diffusion pores, it has been assumed that fluoride permeates Flucs in its charged state, i.e., F^−^ (16). However, as the permeation energetics has not been established, it is unclear if and how Flucs could compensate for the high desolvation penalty of F^−^ (111 kcal/mol, Ref. (27)) upon its entry. A eukaryotic Fluc from *Camellia sinensis* has been shown to couple H^+^ gradient with fluoride efflux (28), but the same coupling has not yet been reported in its bacterial counterparts. The fact that the identity of the “polar track” residues varies among Flucs adds to the mystery. Fluc-Ec2 comprises an ionizable “track” residue H106 (**Fig. 1C**). Replacement of H106, even by HBond substitutes like H106S (Fluc-Bpe equivalence), makes Fluc-Ec2 completely inactive (11), suggesting that H106 is more than an HBond donor. Since H106 lies near the pore end and is (at least partially) solvated, it is likely to be protonated under physiological pHs to coordinate with F83 via cation-π interactions (**Fig. 1BD**). Titration of H106 or F^−^ might be necessary for F^−^ bypass to avoid electrostatic trapping. The absence of an ionizable counterpart in other Flucs would suggest either no titration of H106/F^−^ in Fluc-Ec2 or wide variance in the permeation mechanism across the family. Other “track” residues in Fluc-Ec2 (S102/S110/T114) are insensitive to mutagenesis (11). The lack of necessity of HBond donors at these locations with conserved polarity thus adds more complexity. These mechanical puzzles require a comprehensive assessment of the charge state of fluoride inside Fluc and the accompanying permeation energetics.

Electronic polarization may be essential during ion permeation through some inhomogeneous membrane systems (29,30). This is one major challenge for computational studies using standard nonpolarizable additive force fields (FFs), which approximate the response to different dielectric media in a mean-field fashion (31,32). Additive FFs have, for example, overestimated the transfer barrier of cations (Li^+^/Na^+^/K^+^) through gramicidin A (gA) channel by 5 – 10 kcal/mol, and estimated conductance that differed by several orders of magnitude from experimental values (33-36). Zhang *et al*. (37) noticed that the water wire in gA was more structured with the additive FF description. Ngo *et al*. (38) further confirmed this FF-dependent water structure. They also revealed the balance between water–water and water–protein interactions impacted the water dynamics inside the channel and changed the energetics of ion permeation. Similar observations have also been reported for potassium (KcsA) and voltage-gated sodium (Na_V_) channels. Jing *et al*. (39) discovered polarization played a crucial role in the thermodynamic stabilities of different K^+^ binding configurations. Sun *et al*. (40) found the free energy barrier of Na^+^ transfer was overestimated by ∼2 kcal/mol using additive FF due to a failure to describe the electrostatics between the charged gating residues and aromatic residues with large polarizability. The anion case has, however, received much less attention. Vergara-Jaque *et al*. (41) examined the free energy of I^−^ transfer through a sodium-iodide symporter (NIS) and noticed an additive FF missed a global minimum around the putative I^−^ binding site. However, their work studied a fragment of NIS and did not inspect the origin of the FF-dependent energetics. Fluc, a small channel, should thus be an ideal model for assessing the electronic polarization effect during anion permeation.

In this paper, we have investigated the fluoride permeation mechanism in Fluc-Ec2 via a combination of *in silico* approaches. We first applied the continuous constant-pH molecular dynamics (CpHMD) technique (42,43) to examine the charge state of fluoride inside Fluc. In CpHMD, a titration coordinate, *λ*, is associated with each ionizable site and propagates simultaneously with the spatial coordinates in the MD. The charge states (or *λ* values) are updated on-the-fly during the simulation and closely coupled to the conformational dynamics. With this approach, we found that F^−^ is stable inside Fluc using the membrane-enabled hybrid-solvent CpHMD (44,45), a formulation that provides high-accuracy p*K*_a_ prediction (46). We then evaluated the free energy of F^−^ permeation through Fluc with the replica-exchange umbrella sampling (REUS) method (47,48). We discovered that F^−^ is relayed inside a network of HBonds, ion-pairs, and anion-π pairs. The dehydration penalty of anhydrous S_cen_ F^−^ was mainly offset by the HBonds, suggesting S_cen_ is the selectivity filter discriminating F^−^ over Cl^-^. The selectivity over cations is likely conferred by the “Coulombic plug” R19. Moreover, proper treatment of ion-π interactions requires explicit treatment of the ion-induced polarization of the π-electron cloud, which is not well represented in non-polarizable additive FFs (49-51). Thus, we employed both the CHARMM additive CHARMM36m (52-54) and Drude polarizable (55-61) FFs to draw comparisons. The results showed that the additive FF significantly overestimates the ion-protein interactions, leading to an estimated F^−^ flux about six orders of magnitude slower than experimental measurements. Predictions with the polarizable FF, on the other hand, agree well with the experiments, which underlines the importance of electronic polarization during F^−^ permeation in this channel and may point to the utility of polarizable models in studying anion permeation through membrane proteins in general.

## METHODS

The study was conducted following a protocol previously established (62,63) with necessary modifications. For simulations using additive FF, protein, lipids, and waters were represented by CHARMM36m (53,54), CHARMM36 (52), CHARMM-modified (64) TIP3P (65) models, respectively. The parameters for F^−^ and hydrofluoric acid (HF) were reported by Senn *et al*. (66) and Laage *et al*. (67), respectively. For CpHMD simulations, the protein was modeled by CHARMM22/CMAP (68,69). In the simulations using polarizable FF, the CHARMM Drude-2013 models (55-60) with cation-π and anion-ring corrections (61) were used. The initial conformation of Fluc-Ec2 was retrieved from the crystal structure (PDB: 5A43) (16) and embedded in a 1-palmitoyl-2-oleoyl-*sn*-glycero-3-phosphoethanolamine (POPE) lipid bilayer. The system was built using CHARMM-GUI *Membrane Builder* (70-74) and equilibrated using GROMACS (75) (version 2019.3) for 100 ns. The last snapshot from the equilibration was then converted to CpHMD-compatible format using CHARMM (76) (version c42b2). The membrane-enabled hybrid-solvent CpHMD (44,45) with the pH-based replica-exchange enhanced-sampling protocol (pH-REX) (44) ran for 20 ns per replica using CHARMM to calculate the p*K*_a_s of fluoride and ionizable residues inside Fluc-Ec2. Twenty-one replicates of the CpHMD system were also generated with the fluoride restrained at various locations along the membrane normal (Z-axis) and equilibrated at pH 7 for 50 ns. The equilibrated CpHMD replicates were converted to fixed-protonation-state style using CHARMM for additive and polarizable REUS (48) simulations. The potentials of mean force (PMFs) of F^−^ permeation through Fluc-Ec2 were derived for both sets of systems using NAMD (77). Details of simulations and analysis protocols are described in the Supporting Material.

## RESULTS AND DISCUSSION

### Fluoride is charged inside Fluc-Ec2

We first investigated the charge state of fluoride, i.e., HF or F^−^, inside Fluc-Ec2 using the membrane-enabled hybrid-solvent CpHMD combined with pH-REX (44,45). All Asp, Glu, and His residues were allowed to ionize in the simulation. Na^+^–chelating R19s were also set to be ionizable to evaluate if they stay charged. The pH-REX CpHMD simulation was 20 ns per replica. Calculated p*K*_a_s converged within 10 ns (**Fig. S1**). Final p*K*_a_s are listed in **Table 1**. p*K*_a_s for H60s, D62s, and E86s are shifted from their corresponding model compound values (6.5, 3.7, 4.3) (78) by less than 1 pH units. The “track” residue H106s have significantly upshifted p*K*_a_s (8.6 and 9.6 for chain A and B, respectively), indicating the doubly protonated form is stable around physiological pH. Fluorides and the Na^+^-sandwiching R19s stay deprotonated and protonated, respectively, within the pH range studied (3.5–10.5). The p*K*_a_s suggest that at pH 7, where the Fluc-Ec2 F^−^ efflux was measured (6,10,11), fluoride and H106s should stay charged. During the simulation, F^−^ ions remained around the S_cen_ sites and no F^−^ release was observed, possibly due to limited sampling and overstabilized ion–protein interactions under the additive FF representation (see discussion below).

**TABLE 1.** Calculated p*K*_a_s for ionizable residues/ligands in Fluc-Ec2

| Residue | $pK_a$ | |
| --- | --- | --- |
|  | Chain A | Chain B |
| **R19** | Protonated |  |
| **H60** | 7.3 | 7.4 |
| **D62** | 3.4 | 3.7 |
| **E86** | 3.9 | 3.8 |
| **H106** | 8.6 | 9.6 |
| **HF** | Deprotonated |  |

### F^−^ permeation through Fluc-Ec2 is feasible

Based on the charge states determined by the pH-REX CpHMD, we then employed the REUS technique (47,48) to evaluate the free energy of F^−^ permeation through Fluc-Ec2 using either the C36m (52-54) or Drude (55-61) FFs. Since bacterial Fluc monomers function independently (10), we only modeled F^−^ permeation through monomer A while restraining the F^−^ in monomer B around its crystallographic location. **Fig. 2A** displays the resulting permeation PMFs as a function of the distance from the Fluc center (ΔZCOM), which delineates the change in free energy as F^−^ travels between internal and external aqueous bulks via Fluc. Both FFs give PMFs (black for C36m, red for Drude) that agree on their overall trend: a rugged free energy landscape with a global minimum close to the S_cen_ binding site (ΔZCOM of – 4 Å). However, the absolute values of the PMFs differ significantly so that F^−^ is much more stable under the C36m representation. In C36m, F^−^ needs to cross a free energy barrier of 17.7 ± 0.8 kcal/mol, but in Drude, the barrier is *ca*. 60% smaller (7.4 ± 0.3 kcal/mol). Moreover, the S_cen_ minimum is only 0.6 ± 0.5 kcal/mol more stable than the S_ext_ one (ΔZCOM of 4 Å) in the polarizable Drude FF. This suggests that, while slightly favored at S_cen_, F^−^ is also likely to be found at S_ext_. In addition, a local minimum at S_int_ was also found.

**FIGURE 2.**
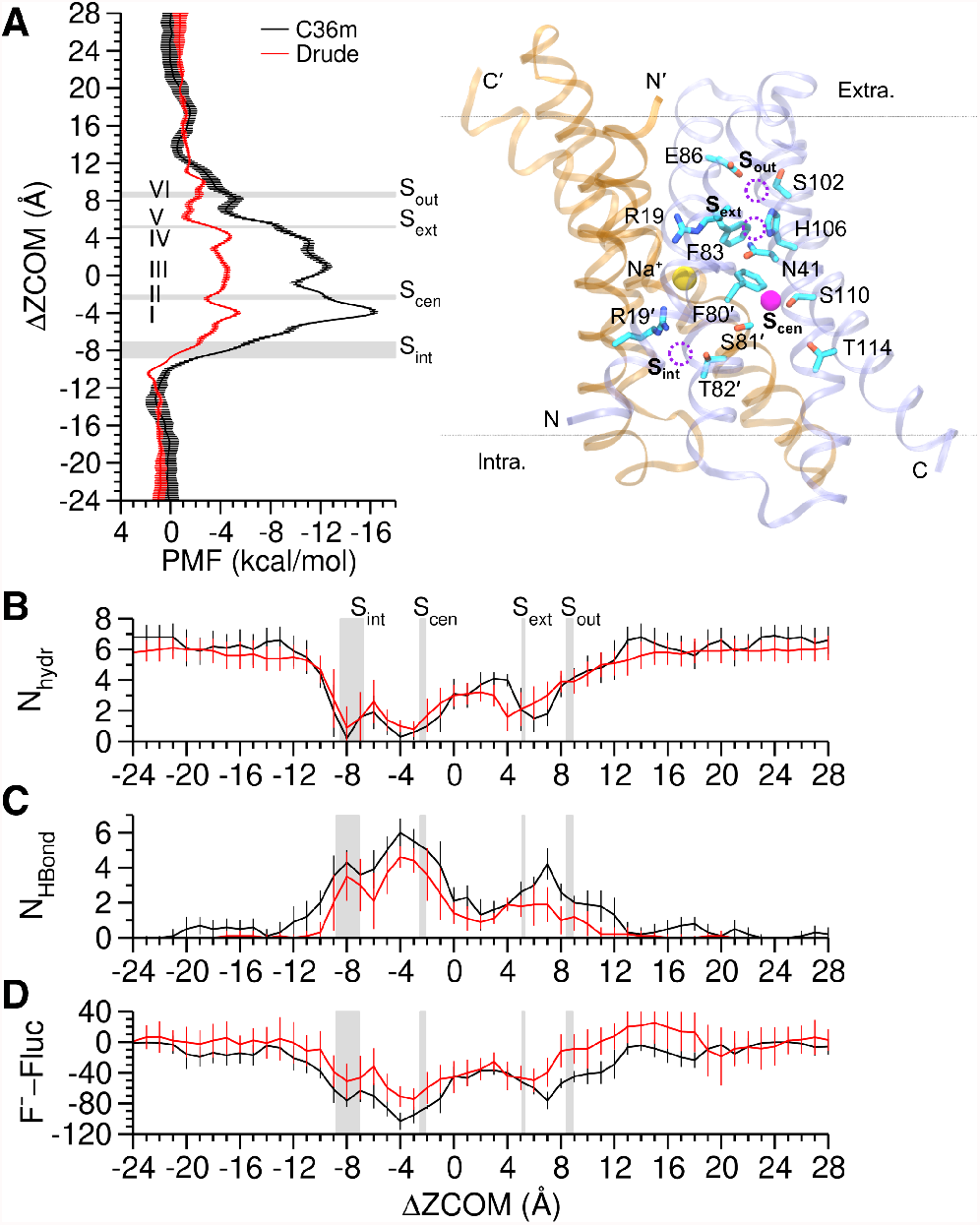
F^−^ permeation thermodynamics. **(A)** Potential of mean force (PMF) as a function of ΔZCOM, which is defined as the center-of-mass (COM) distance between F^−^ and the Fluc Cα-atoms projected on the bilayer normal (Z-axis). PMFs from the additive C36m and polarizable Drude FFs were set to zero in aqueous bulk. Error bars are the standard deviations calculated from block analysis. Grey boxes indicate the crystallographic locations of S_int_, S_cen_, S_ext_, and S_out_ F^−^ (centroid and halved box width respectively represent the mean and standard deviation calculated from all available PDB structures). Roman numerals I–VI indicates the representative states illustrated in **Fig. 3**. The Fluc-Ec2 structure, with essential structural features labeled, is aligned with the PMFs. Dashed purple circles indicate S_int_, S_ext_, and S_out_. **(B)** Hydration number of F^−^, defined as the number of water oxygen atoms within 3.5 Å of F^−^. **(C)** Number of HBonds between F^−^ and Fluc. HBond is considered to be present if the donor–H distance is ≤ 2.4 Å. **(D)** Interaction energy (kcal/mol) of F^−^ with Fluc. The means and fluctuations (shown as error bars) were calculated every 1 Å along ΔZCOM.

To better understand the permeation energetics, we quantified the hydration number (N_hydr_) and the number of HBonds (N_HBond_) of F^−^ as a function of ΔZCOM. As displayed in **Fig. 2B**, F^−^ is dehydrated inside Fluc. Yet, the desolvation penalty is compensated by HBonds with Fluc (**Fig. 2C**), as the N_hydr_ and N_HBond_ profiles are complementary, regardless of FFs. For instance, the anhydrous S_cen_ F^−^ is stabilized by 6 HBonds. The S_ext_ F^−^ is solvated by at least 2 water molecules and fewer HBonded. Since N_HBond_ excludes water, our results indicate F^−^ is constantly coordinated with Fluc and water molecules via ∼ 6 HBonds during the permeation, which approximately equals the N_hydr_ in aqueous bulk (**Fig. 2B**). Following the protocol by Lin *et al*. (79), we calculated the interaction energy (E_inter_) between F^−^ and Fluc (including the central Na^+^ and the other F^−^). Consistent with the PMFs, both FFs predict overall favorable E_inter_ of F^−^ with Fluc, but E_inter_ is stronger in C36m (**Fig. 2D**). Residue breakdown of E_inter_ indicates that the C36m overstabilization mainly originates from the interaction with R19′ around S_int_ and S_cen_, and with R19 around S_out_, which are overestimated by 15 – 20 kcal/mol (**Fig. S2**). Moreover, C36m predicts E_inter_ constantly favorable throughout Fluc, but Drude predicts unfavorable E_inter_ around S_out_ and beyond(ΔZCOM > 8 Å, **Fig. 2D**), where the negatively charged E86 resides (**Fig. 2A, Fig. S3A**). Because F^−^ is more likely to form contact with E86 in Drude than in C36m (**Fig. S3B**). Decomposition of E_inter_ indicates unfavorable self-polarization energy (E_self_), which quantifies the contribution of electronic polarization to E_inter_ in Drude and is absent in C36m (79), dominates F^−^–E86 (**Fig. S3C**). Unfavorable Drude E_inter_, though smaller, is also seen with the “Phe-box” (**Fig. S2**) due to unfavorable E_self_ (**Fig. S4**). By contrast, the interactions are marginal in C36m (**Fig. S2**). The only exception is F80′, which strongly interacts with the S_cen_ F^−^ via HBond (**Fig. 3B**).

**FIGURE 3.**
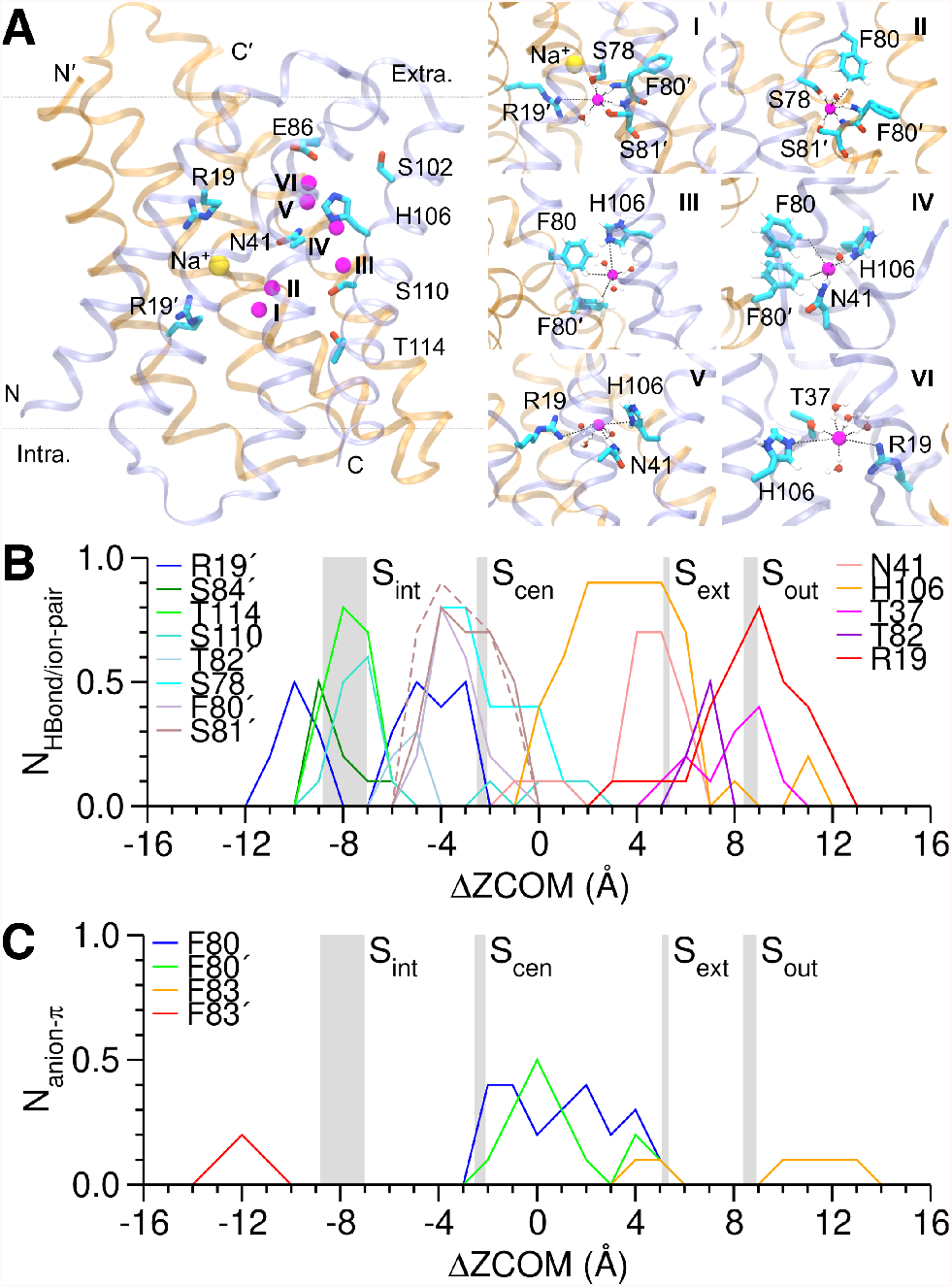
F^−^ permeation mechanism. **(A)** Representative conformations during the permeation. Roman numerals are consistent with **Fig. 2A**. The relative positions of I–VI are indicated in a full-length Fluc-Ec2. **(B)** Occupancies of HBonds/ion-pairs between F^−^ and Fluc. HBond is considered to form if the donor–acceptor distance is ≤ 3.5 Å and the donor–H–acceptor angle is ≥ 150°. Ion-pair is considered to form if the N–F^−^ minimal distance is ≤ 4.0 Å. Note S81′ HBonds with F^−^ via backbone (solid line) and side-chain (dashed line). **(C)** Occupancies of anion-π pairs between F^−^ and the “Phe-box”. Anion-π pair is present if F^−^ is within 4.5 Å of any ring carbon atoms and subtends an angle of ≤ 35° with the ring plane. For clarity, error bars (fluctuations) are not shown. All data were extracted from the Drude REUS simulations.

We next examined the permeation kinetics. The local diffusion constant *D* was calculated to be (2.1 ± 0.1) × 10^-5^ and (1.4 ± 0.2) × 10^-5^ cm^2^/s respectively from the C36m and Drude REUS simulations (**Table 2**) using the Woolf-Roux equation (80). The experimental *D* is 1.475 × 10^-5^ cm^2^/s in infinitely dilute aqueous solution (81), and (1.4 ± 0.1) × 10^-5^ cm^2^/s considering finite concentration (82). Our results therefore indicate that the Drude FF predicts a correct *D* whereas C36m overestimates *D* by 50%, which is consistent with the study of ion conductivity by Prajapati *et al*. (83). The permeation rate constant *k* was calculated from the PMF and *D* (**Eqs. S2-S3**) based on the mean first passage time model (84). The resulting *k* is (4 ± 1) × 10^6^ and (0.29 ± 0.26) ions/s for Drude and C36m, respectively (**Table 2**). The Drude estimate agrees with the experimental turnover rate of 10^6^–10^7^ ions/s (6,8). But the C36m estimate is six orders of magnitude slower, far below the typical rate for ion channels. These results further support the viability of F^−^ permeation through Fluc and point to nonnegligible electronic polarization during permeation.

**TABLE 2.** F^−^ bulk diffusion constant *D* and permeation rate constant *k*

| | $D (10^{-5} \text{ cm}^2/\text{s})$ | $k (\text{ions/s})$ |
| --- | --- | --- |
| **Drude** | $1.4 \pm 0.2$ | $(4 \pm 1) \times 10^6$ |
| **C36m** | $2.1 \pm 0.1$ | $0.29 \pm 0.26$ |
| **Experiment** | $1.4 \pm 0.1$ | $10^6$–$10^7$ |

### F^−^ relays within a network of non-bonded interactions

Finally, we investigated the atomistic permeation mechanism based on the Drude simulations. We first computed the occupancies of HBonds and ion-pairs between F^−^ and Fluc polar residues (**Fig. 3B**) as well as anion-π pairs with the “Phe-box” (**Fig. 3C**) along the pore (i.e., ΔZCOM). We then compared these interaction profiles with the permeation PMF (**Fig. 2A**) to extract the most representative patterns (**Fig. 3A**) for metastable and transition states I–VI (**Fig. 2A**).

In the permeation direction (**Fig. 2A**), F^−^ enters Fluc and binds to S_int_ with a sharp decrease in N_hydr_ (ΔZCOM from –11 to –5 Å, **Fig. 2B**), but the interactions with R19′, S84′, T114, and S110 (**Fig. 3B**) stabilize the desolvated F^−^. Note S_int_ is not a minimum in the permeation PMF (**Fig. 2A**), which explains why S_int_ was not captured by the structures crystallized with F^−^. The F^−^ anion translocates from S_int_ to S_cen_ while interacting with R19′, S78, F80′, S81′, and T82 (**Fig. 3B**). Once bound to S_cen_ (state I, ΔZCOM from –5 to –3 Å), F^−^ predominantly forms 5 HBonds with S78, F80′, S81′ (backbone and side-chain), and a water molecule. Meanwhile, F^−^ ion-pairs with R19′ (**Fig. 3AB**). As a result, despite the lower solvation (**Fig. 2B**), F^−^ is stabilized by the most abundant HBonds and ion-pairs in state I (**Fig. 2C**), leading to the global minimum in the PMF (**Fig. 2A**). Note that the F^−^–Na^+^ pair is not close enough to be considered as an ion-pair. State II (ΔZCOM from –3 to –1 Å) is the first transition state. Compared to state I, the ion-pair with R19′ is lost in state II, while the 5 HBonds retain (**Fig. 3AB**). F^−^ moves to the proximity of F80 to form an anion-π pair (**Fig. 3AC**), consistent with the crystal S_cen_ binding pattern. The anion-π pair is not strong enough to balance the ion-pair loss, which explains why state II is less stable. The HBond with S110 (S112 in Fluc-Bpe, **Fig. 1B**) was also observed in our simulations but with marginal occupancy (**Fig. 3B**). In state III (ΔZCOM from –1 to 3 Å), F^−^ becomes the most solvated (**Fig. 2B**) but the least HBonded (**Fig. 2C**). With all the HBonds in states I and II absent, F^−^ begins ion-pairing with H106 (**Fig. 3AB**). Additional stabilization is provided by anion-π pairs with F80 and F80′ (**Fig. 3AC**) and occasionally by HBond with N41 (**Fig. 3B**), making state III a local minimum in the PMF (**Fig. 2A**). State IV (ΔZCOM from 3 to 5 Å) is another local minimum in the PMF and corresponds to S_ext_ (**Fig. 2A**). State IV differs from state III by less solvation (**Fig. 2B**) and stronger HBond with N41 (**Fig. 3AB**). The move from state IV to state V (ΔZCOM from 5 to 7 Å), the second transition state, is accompanied by gradual loss of the interactions with F80s and N41 (**Fig. 3BC**) and by increasing hydration (**Fig. 2B**). Starting from state V, F^−^ ion-pairs with R19 (**Fig. 3AB**). State VI (ΔZCOM from 7 to 11 Å) matches S_out_ and is the last local minimum in the PMF (**Fig. 2A**), where the solvated F^−^ is coordinated mainly with R19 and T37 (**Fig. 3AB**) and infrequently with F83 (**Fig. 3C**).

Our results show that F^−^ permeates by relaying inside a non-bonded network. The identified interaction patterns highlight the indispensability of conserved or immutable residues. Most of the Fluc interior is an HBond “desert” except around S_cen_ (**Fig. 2C**). Thus, extra stabilization is required to offset the desolvation penalty and help F^−^ permeate. First, beyond S_cen_ (ΔZCOM > –1 Å), F^−^ primarily HBonds with N41 and ion-pairs with H106 (ΔZCOM from 2 to 6 Å, **Fig. 3B**). Within the “gap”, the anion-π pairs with F80s govern (**Fig. 3C**), consistent with the observation that F80 mutants are nonfunctional (10,16). Next, N41 and H106 stabilize and guide F^−^ to S_ext_ (**Fig. 3B**). Our CpHMD simulation revealed that the F^−^–H106 ion-pair facilitates the permeation (**Fig. S5**). This helps to explain the puzzling finding that H106 is irreplaceable (11). Then, the Drude PMF predicts S_cen_ as a transition state (**Fig. 2A**), which appears to be a discrepancy as the crystal binding site should be a minimum in PMF. However, most Fluc structures were crystallized around pH 6 and below (**Table S1**), whereas our simulations modeled pH 7. Given the stronger F^−^–H106 ion-pair at acidic pHs, it is reasonable the crystal S_cen_ site upshifts relative to our PMF global minimum (**Fig. S5**). Lastly, two R19s are found near the two pore openings (**Fig. 1A**). They likely help retrieve F^−^ from internal/external bulk regions and provide initial stabilization upon pore entry (**Fig. 3B**). This scenario also holds for F83s via anion-π pairs, although the interactions are weaker (**Fig. 3C**).

## CONCLUDING DISCUSSION

We have studied the mechanism of fluoride permeation through Fluc-Ec2 *in silico*. CpHMD simulation found a stable F^−^ (**Table 1**) at the most dehydrated S_cen_ site (**Fig. 2B**). The permeation energetics at pH 7 were then established by the REUS free energy calculations. The permeation PMF from the electronically polarizable Drude simulations revealed a global minimum around S_cen_, a slightly less stable local minimum at S_ext_, and a third local minimum at S_out_ (**Fig. 2A**), which are consistent with the crystal structures (**Table S1**). F^−^ is dehydrated during the permeation (**Fig. 2B**), but the desolvation penalty is compensated by HBonds within Fluc (**Fig. 2C**) so that F^−^ stays as coordinated inside the channel as it is in aqueous bulk. The permeation rate estimated from the Drude simulations agrees well with the experimental turnover rate measured at pH 7 (6,8), further supporting that Fluc transfers F^−^ anion. We note, however, that the 1-dimensional permeation PMFs (**Fig. 2A**) are single-ion PMFs which biased the position of a permeating F^−^ along the pore regardless of if and how other F^−^ ions might simultaneously bind. We are unable to conclude at this stage if the “knock-on” mechanism in potassium channels (85,86), in which one incoming ion collides and pushes other ions through the pore, also applies to Fluc, as suggested by McIlwain *et al*. (19). Elucidating stable multi-ion binding patterns necessitates very demanding multi-dimensional free energy calculations which take into account the relative positions of multiple ions as other collective variables (39,86-89).

F^−^ has been proposed to be stabilized by non-bonded interactions with the pore-facing residues during the permeation (16). Our study supports this idea by showing that F^−^ relays inside a non-bonded network and also explains residue conservation/indispensability. R19 acts as an “antenna” for fetching F^−^ from aqueous bulks and, more importantly, stabilizes F^−^ to S_cen_ or S_ext_ (**Fig. 3AB**). Though not yet reported in bacterial Flucs, mutating its equivalent in yeast *S. cerevisiae* (FEX_Sc_) significantly reduces F^−^ efflux (8). N41 has further been shown in both bacteria (16) and eukaryotes (8) to be an immutable HBond donor, with side-chain rotation thought to be coupled with F^−^ motion (16). We observed that the HBond with N41 guided the S_cen_ F^−^ to S_ext_ (**Fig. 3AB**), supporting the “rotameric switch” conjecture. Polar residues S78, S81, T82, and S84 (T80, S83, T84, and S86 in Fluc-Bpe) were also identified as HBond donors in our simulations (**Fig. 3B**). The residues S78 and S81 form strong HBonds with the S_cen_ F^−^. T82 and T82′ help F^−^ travel from S_int_ to S_cen_ and from S_ext_ to S_out_, respectively. These results are consistent with the mutagenesis that S81A and S81A/T82A respectively cause ∼50% and 100-fold conductance loss (19). It is unclear, however, why F^−^ efflux was completely abolished in S81T (19). We speculate that –CH_3_ is so bulky at S_cen_ that it eliminates more HBonds than the one with the S81 side-chain (e.g., the water HBond, **Fig. 3A**), making F^−^ highly unstable here. S84 HBonds with the S_cen_–marching/leaving F^−^. The irreplaceable “Phe-box” (10,11,16) has been hypothesized to form edgewise anion-π pairs with F^−^ (16), which was confirmed by our simulations (**Fig. 3C**). Notably, we found the anion-π pairs with F80s were prominent during the transition between S_cen_ and S_ext_. F80 also HBonds with the S_cen_ F^−^ (**Fig. 3B**). The anion-π pairs with F83s seen near the pore ends are considerably weaker (**Fig. 3C**). The variation in their pairing abilities is consistent with the observation that Fluc is resistant to F80M rather than F83M (11), because electropositive Met is an anion-π substitute for Phe. However, the irreplaceability of F83 remains mysterious.

The “polar track” (**Fig. 1D**) was proposed to HBond F^−^ (16), but this argument was challenged by a follow-up mutagenesis study disclosing several paradoxes (11). First, mutations S102A and S110A do not impair Fluc function. No HBond with S102 was seen in our simulations. F^−^ HBonds with S84′, T114, and S110 in the same region (**Fig. 3B**), so removing S110 is likely not deleterious. Second, mutants T114S and T114A are completely inactive, but aromatic substitutes such as T114V and T114I are fully active. These results indicate no HBond is necessary at this location. To reach S_cen_, the S_int_ F^−^ has to pass a hydrophobic gate formed by T114, I48, L52, V85′, and F88′. T114S may not be bulky enough to preserve gate integrity. We speculate the HBonds with S110 or S84′ could help stabilize the dehydrating F^−^ passing the gate (ΔZCOM of –8 Å, **Fig. 2B**). Although Fluc is insensitive to mutations S84A or S110A (8,11), it would be interesting to test *in vitro* if the double mutant S84A/S110A is active. Third, H106 is immutable in Fluc-Ec2 as neither aromatic nor polar substitution is tolerated. H106 plays the same role as N41 via an ion-pair with F^−^ (**Fig. 3B**) in our simulations, which seemingly explains its irreplaceability, but it is unknown why H106 is not strictly conserved. S102 and H106 are respectively present as Y104 and S108 in Fluc-Bpe (**Fig. 1D**), a pattern shared by FEX_Sc_ (Y345 and S349). It is surprising that FEX_Sc_ tolerates mutations S349A, S349V, and Y345F but is highly sensitive to Y345A (12). In Fluc-Bpe, Y104 only permits the conservative mutation Y104F, but not replacements by Ser, His, or Ile (19). As F^−^ likely coordinates a Tyr side-chain via both anion-π pair and hydroxyl HBond, these results imply that if the interactions with residue 106 (Fluc-Ec2 numbering) are reduced, stabilization from residue 102 becomes essential. In other words, the conserved polarity requires precise geometrical side-chain arrangement. Furthermore, two strongly conserved residues, T37 (T39 in Fluc-Bpe) and E86, are found in proximity to H106. In Fluc-Bpe, T39 is replaceable by Ser, but not by Ala, Asn, Cys, or Val, whereas E88 tolerates Ala, Asp, and Gln but not Lys (19). Mutating their FEX_Sc_ equivalences causes severe function loss (12).T37 is the sole stable HBond donor around S_out_ (**Fig. 3B**). Crystal structures capture F^−^ coordination with E86 (**Fig. S3A**), but our simulations found no close contact between the two (**Fig. S3B**) because both were charged. A recent study found that E86 does not ionize between pH 7 and 8.7 and suggests that E86 contributes to anion recognition (19). CpHMD predicted an E86 p*K*_a_ that accords with the experiment (∼4, **Table 1**). To avoid electrostatic repulsion, either E86 or the permeating F^−^ ion should (transiently) protonate likely by accepting a proton from the surrounding water molecules (**Fig. 2B**) so that the S_out_ F^−^ and E86 could form close contact. Note that X-ray diffraction cannot distinguish isoelectronic fluoride and water, the electron density at S_out_ was thus assigned as water molecules without further evidence of anionic character in many crystal structures (16). Rigorous evaluation of the fluoride stability at S_out_ requires quantum mechanical modeling, which may also aid in clarifying the role of E86. Although the possibility of protonation on either E86 or F^−^cannot be readily excluded at this stage and may be the focus of future work, it should not negate the major conclusion of the current work; that is, Fluc transfers F^−^.Our study also offers atomistic insight into the remarkable selectivity for F^−^. The R19 residue generates a positive “Coulombic plug” at either end of the pore and explains the selectivity for F^−^ over Na^+^ (6). This is supported by the fact that replacing the Arg by Lys has little impact on FEX_Sc_ growth (8). The same feature is seen in a cationic transporter EmrE which comprises a conserved E14 at a spatially similar location (90,91). Although pore size may also be a factor, the interaction patterns during the permeation (**Fig. 3AB**) suggest that S_cen_ is more likely to be the selectivity filter for F^−^ over Cl^−^ (6,8). The F^−^ anion is mainly stabilized by HBonds (**Fig. 3AB**) in this anhydrous (**Fig. 2B**) region. Considering the weaker HBonding capability of Cl^−^ than that of F^−^ (92,93), it should be improbable for Cl^−^ to offset the desolvation penalty (81 kcal/mol, Ref. (27)) at S_cen_. McIlwain *et al*. (19) recently examined the competitive binding of several halides and pseudohalides to S_out_ and discovered that the anion’s ability of accepting a HBond contributes to its recognition by Flucs. Our study lends credence to the notion that HBond is the key to the selectivity of Flucs among anions. However, we can see from the different desolvation penalties of F^−^ and Cl^-^ that this argument is based on an assumption that the weaker HBonds affect the channel more than desolvation. We are unable to perform a quantitative comparison of these two effects from our 1-dimensional PMFs for F^−^ permeation. To do so requires demanding 2-dimension free energy calculations that bias solvation as a second collective variable (94-97) for both anions.

Another extraordinary feature of Fluc is the Na^+^ buried at the dimer interface (**Fig. 1A**) and liganded with four backbone carbonyl oxygen atoms from both monomers (G75s and S78s, **Fig. S6A**; G77s and T80s in Fluc-Bpe). These Na^+^–O=C pairs were quite stable in our simulations (**Fig. S6B**), making Na^+^ a cation that may help keep the two monomers closely “glued”. The same observation was reported by Ariz-Extreme *et al*. (98). In addition, it was curious that Na^+^ is tetrahedrally coordinated in Fluc crystal structures (5,14,16), as Na^+^ has a typical coordination number (N_coord_) of 5 – 6. We found the Na^+^ was always liganded with one water molecule (**Fig. S6C**) and occasionally with another water molecule, the carbonyl oxygens of G76s, or the side-chain hydroxyl oxygens of S78s, leading to a stable N_coord_ of 5 – 6 (**Fig. S6D**). Moreover, the Na^+^ is located on the two-fold symmetry axis parallel to the membrane surface (16). Water, being the fifth stable ligand coordinating the Na^+^, breaks the symmetry of the overall structure. Electronically polarizable models can provide a more physically realistic representation for simulating ion transport across heterogeneous membrane systems with complex dielectric environments (29-32). In this study, we have compared the nonpolarizable additive C36m and polarizable Drude-2013 FFs. A significantly overestimated permeation barrier around S_cen_ (**Fig. 2A**) and consequently a greatly underestimated flux (**Table 2**) were given by the additive simulations, consistent with existing computational studies in cation channels (33-36,38-40). A breakdown by residue of the F^−^–Fluc E_inter_ indicates the additive overstabilization primarily originates from the Coulombic interactions with R19s, which was 10 – 20 kcal/mol stronger in C36m (**Fig. S2, S7-S9**). It is not surprising to see more substantial charge–charge interactions in additive FFs because dielectric constants are underestimated (31,32). However, no large difference was found for the interactions with H106 (**Fig. S2, S7-S9**). In the case of interactions with π electron-rich residues such as F80s, F83s or charged E86s, C36m captured little interaction, but Drude yielded interactions of 5 – 20 kcal/mol (**Fig. S2**) dominantly contributed by E_self_ (**Fig. S3-S4**), consistent with previous studies (98,99). This difference is due to a limited description of the ion-induced polarization of π electron in additive FFs (61), which E_self_ accounts for in Drude FFs (31,79). Because F80′ HBonds with the S_cen_ F^−^ (**Fig. 3AB**), E_elec_ is much stronger around S_cen_ (**Fig. S4**). But E_self_ dominates when F^−^ transits between S_cen_ and S_ext_ (**Fig. S4**). During the transit, polar residue N41 also experiences an E_self_ of up to 10 kcal/mol (**Fig. S9**), but E_elec_ is essential (**Fig. S2, S8**) due to the HBond with F^−^ (**Fig. 3AB**). Furthermore, while the force field performance was assessed using single-ion PMFs, a recent KcsA study considering the multi-ion effect has shown that electronic polarization is critical for K^+^ (39). Given the higher polarizability of F^−^ (56,57), we argue that including a study of the multi-ion effect would further support the notion that electronic polarization should be considered in anion channels/transporters.

In summary, we have computationally investigated the fluoride permeation mechanism through Fluc-Ec2. We confirm F^−^ permeation with a calculated rate in encouraging agreement with experiment and we reveal that the permeation is enabled by F^−^ relaying inside a non-bonded network. Although the variance in the pore-lining polar residues remains somewhat puzzling, the present work paves the way for future studies to elucidate delicate control of the pore polarity for rapid F^−^ transit. Lastly, this work highlights the possible importance of electronic polarization during anion permeation through membrane channels/transporters and the necessity of using polarizable models for treating such processes.

## SUPPORTING MATERIAL

Supporting material can be found online at https://doi.org/10.1016/j.bpj.xxxx.xx.xxx.

## AUTHOR CONTRIBUTIONS

G.A.V. designed the research. Z.W. and Z.Y. ran the simulations. Z.W., Z.Y., and G.A.V. analyzed the data. Z.W., Z.Y., and G.A.V. wrote the article.

## ACKNOWLEDGMENTS

This research was supported by the National Institute of General Medical Sciences (NIGMS) of the National Institutes of Health (NIH Grant R01 GM053148). The computational resources were provided by the University of Chicago Research Computing Center (RCC). The authors thank Dr. Chenghan Li for his insightful discussion of the implication of permeation PMFs in ion selectivity.

## DECLARATION OF INTEREST

The authors declare no competing interests.

## SUPPORTING CITATIONS

References (100-122) appear in the Supporting material.

## Notes

### Competing Interest Statement

The authors have declared no competing interest.

## REFERENCES

1. Weinstein, L. H., and A. Davison. 2004. Fluorides in the Environment: Effects on Plants and Animals, 1st ed. CABI Publishing, Cambridge, MA.

2. Marquis, R. E. 1995. Antimicrobial actions of fluoride for oral bacteria. Can. J. Microbiol. 41:955–964.

3. Marquis, R. E., S. A. Clock, and M. Mota-Meira. 2003. Fluoride and organic weak acids as modulators of microbial physiology. FEMS Microbiol. Rev. 26:493–510.

4. Stockbridge, R. B., H.-H. Lim, R. Otten, C. Williams, T. Shane, Z. Weinberg, and C. Miller. 2012. Fluoride resistance and transport by riboswitch-controlled CLC antiporters. Proc. Natl. Acad. Sci. U.S.A. 109:15289– 15294.

5. Macdonald, C. B., and R. B. Stockbridge. 2017. A topologically diverse family of fluoride channels. Curr. Opin. Struct. Biol. 45:142–149.

6. Stockbridge, R. B., J. L. Robertson, L. Kolmakova-Partensky, and C. Miller. 2013. A family of fluoride-specific ion channels with dual-topology architecture. eLife 2:e01084.

7. Ji, C. H., R. B. Stockbridge, and C. Miller. 2014. Bacterial fluoride resistance, Fluc channels, and the weak acid accumulation effect. J. Gen. Physiol. 144:257–261.

8. Smith, K. D., P. B. Gordon, A. Rivetta, K. E. Allen, T. Berbasova, C. Slayman, and S. A. Strobel. 2015. Yeast Fex1p Is a Constitutively Expressed Fluoride Channel with Functional Asymmetry of Its Two Homologous Domains. J. Biol. Chem. 290:19874–19887.

9. Turman, D. L., J. T. Nathanson, R. B. Stockbridge, T. O. Street, and C. Miller. 2015. Two-sided block of a dual-topology F– channel. Proc. Natl. Acad. Sci. U.S.A. 112:5697–5701.

10. Last, N. B., L. Kolmakova-Partensky, T. Shane, and C. Miller. 2016. Mechanistic signs of double-barreled structure in a fluoride ion channel. eLife 5:e18767.

11. Last, N. B., S. Sun, M. C. Pham, and C. Miller. 2017. Molecular determinants of permeation in a fluoride-specific ion channel. eLife 6:e31259.

12. Berbasova, T., S. Nallur, T. Sells, K. D. Smith, P. B. Gordon, S. L. Tausta, and S. A. Strobel. 2017. Fluoride export (FEX) proteins from fungi, plants and animals are ‘single barreled’ channels containing one functional and one vestigial ion pore. PLoS ONE 12:e0177096.

13. Turman, D. L., and R. B. Stockbridge. 2017. Mechanism of single-and double-sided inhibition of dual topology fluoride channels by synthetic monobodies. J. Gen. Physiol. 149:511–522.

14. McIlwain, B. C., K. Martin, E. A. Hayter, and R. B. Stockbridge. 2020. An Interfacial Sodium Ion is an Essential Structural Feature of Fluc Family Fluoride Channels. J. Mol. Biol. 432:1098–1108.

15. Stockbridge, R. B., A. Koide, C. Miller, and S. Koide. 2014. Proof of dual-topology architecture of Fluc F– channels with monobody blockers. Nat. Commun. 5:5120.

16. Stockbridge, R. B., L. Kolmakova-Partensky, T. Shane, A. Koide, S. Koide, C. Miller, and S. Newstead. 2015. Crystal structures of a double-barrelled fluoride ion channel. Nature 525:548–551.

17. Turman, D. L., A. Z. Cheloff, A. D. Corrado, J. T. Nathanson, and C. Miller. 2018. Molecular Interactions between a Fluoride Ion Channel and Synthetic Protein Blockers. Biochemistry 57:1212–1218.

18. McIlwain, B. C., S. Newstead, and R. B. Stockbridge. 2018. Cork-in-Bottle Occlusion of Fluoride Ion Channels by Crystallization Chaperones. Structure 26:635–639.

19. McIlwain, B. C., R. Gundepudi, B. B. Koff, and R. B. Stockbridge. 2021. The fluoride permeation pathway and anion recognition in Fluc family fluoride channels. eLife 10:e69482.

20. Sand, O., M. Gingras, N. Beck, C. Hall, and N. Trun. 2003. Phenotypic characterization of overexpression or deletion of the Escherichia coli crcA, cspE and crcB genes. Microbiology 149:2107–2117.

21. Rapp, M., E. Granseth, S. Seppälä, and G. von Heijne. 2006. Identification and evolution of dual-topology membrane proteins. Nat. Struct. Mol. Biol. 13:112–116.

22. Weinberg, Z., J. X. Wang, J. Bogue, J. Yang, K. Corbino, R. H. Moy, and R. R. Breaker. 2010. Comparative genomics reveals 104 candidate structured RNAs from bacteria, archaea, and their metagenomes. Genome Biol. 11:R31.

23. Baker, J. L., N. Sudarsan, Z. Weinberg, A. Roth, R. B. Stockbridge, and R. R. Breaker. 2012. Widespread Genetic Switches and Toxicity Resistance Proteins for Fluoride. Science 335:233–235.

24. Li, S., K. D. Smith, J. H. Davis, P. B. Gordon, R. R. Breaker, and S. A. Strobel. 2013. Eukaryotic resistance to fluoride toxicity mediated by a widespread family of fluoride export proteins. Proc. Natl. Acad. Sci. U.S.A. 110:19018–19023.

25. Smart, O. S., J. G. Neduvelil, X. Wang, B. A. Wallace, and M. S. P. Sansom. 1996. HOLE: A program for the analysis of the pore dimensions of ion channel structural models. J. Mol. Graph. 14:354–360.

26. Lomize, M. A., I. D. Pogozheva, H. Joo, H. I. Mosberg, and A. L. Lomize. 2012. OPM database and PPM web server: resources for positioning of proteins in membranes. Nucleic Acids Res. 40:D370–D376.

27. Marcus, Y. 1991. Thermodynamics of Solvation of Ions. Part 5.—Gibbs Free Energy of Hydration at 298.15 K. J. Chem. Soc. Faraday Trans. 87:2995–2999.

28. Song, J. L., C. C. Hou, J. X. Guo, Q. Niu, X. H. Wang, Z. J. Ren, Q. Zhang, C. X. Feng, L. Y. Liu, W. Tian, and L. G. Li. 2020. Two New Members of CsFEXs Couple Proton Gradients to Export Fluoride and Participate in Reducing Fluoride Accumulation in Low-Fluoride Tea Cultivars. J. Agric. Food Chem. 68:8568–8579.

29. Flood, E., C. Boiteux, B. Lev, I. V. Vorobyov, and T. W. Allen. 2019. Atomistic Simulations of Membrane Ion Channel Conduction, Gating, and Modulation. Chem. Rev. 119:7737–7832.

30. Maffeo, C., S. Bhattacharya, J. Yoo, D. Wells, and A. Aksimentiev. 2012. Modeling and Simulation of Ion Channels. Chem. Rev. 112:6250–6284.

31. Lemkul, J. A., J. Huang, B. Roux, and A. D. MacKerell Jr. 2016. An Empirical Polarizable Force Field Based on the Classical Drude Oscillator Model: Development History and Recent Applications. Chem. Rev. 116:4983–5013.

32. Jing, Z., C. Liu, S. Y. Cheng, R. Qi, B. D. Walker, J.-P. Piquemal, and P. Ren. 2019. Polarizable Force Fields for Biomolecular Simulations: Recent Advances and Applications. Annu. Rev. Biophys. 48:371–394.

33. Patel, S., J. E. Davis, and B. A. Bauer. 2009. Exploring Ion Permeation Energetics in Gramicidin A Using Polarizable Charge Equilibration Force Fields. J. Am. Chem. Soc. 131:13890–13891.

34. Vorobyov, I., B. Bekker, and T. W. Allen. 2010. Electrostatics of Deformable Lipid Membranes. Biophys. J. 98:2904–2913.

35. Peng, X. D., Y. B. Zhang, H. Y. Chu, Y. Li, D. L. Zhang, L. R. Cao, and G. H. Li. 2016. Accurate Evaluation of Ion Conductivity of the Gramicidin A Channel Using a Polarizable Force Field without Any Corrections. J. Chem. Theory Comput. 12:2973–2982.

36. Manin, N., M. C. da Silva, I. Zdravkovic, O. Eliseeva, A. Dyshin, O. Yaşar, D. R. Salahub, A. M. Kolker, M. G. Kiselev, and S. Y. Noskov. 2016. LiCl solvation in N-methyl-acetamide (NMA) as a model for understanding Li+ binding to an amide plane. Phys. Chem. Chem. Phys. 18:4191–4200.

37. Zhang, Y., K. Haider, D. Kaur, V. A. Ngo, X. Cai, J. Mao, U. Khaniya, X. Zhu, S. Noskov, T. Lazaridis, and M. R. Gunner. 2021. Characterizing the Water Wire in the Gramicidin Channel Found by Monte Carlo Sampling Using Continuum Electrostatics and in Molecular Dynamics Trajectories with Conventional or Polarizable Force Fields. J. Comput. Biophys. Chem. 20:111–130.

38. Ngo, V., H. Li, A. D. MacKerell Jr, T. W. Allen, B. Roux, and S. Noskov. 2021. Polarization Effects in Water-Mediated Selective Cation Transport across a Narrow Transmembrane Channel. J. Chem. Theory Comput. 17:1726–1741.

39. Jing, Z., J. A. Rackers, L. R. Pratt, C. Liu, S. B. Rempe, and P. Ren. 2021. Thermodynamics of ion binding and occupancy in potassium channels. Chem. Sci. 12:8920–8930.

40. Sun, R.-N., and H. Gong. 2017. Simulating the Activation of Voltage Sensing Domain for a Voltage-Gated Sodium Channel Using Polarizable Force Field. J. Phys. Chem. Lett. 8:901–908.

41. Vergara-Jaque, A., P. Fong, and J. Comer. 2017. Iodide Binding in Sodium-Coupled Cotransporters. J. Chem. Inf. Model. 57:3043–3055.

42. Lee, M. S., F. R. Salsbury Jr., and C. L. Brooks III. 2004. Constant-pH Molecular Dynamics Using Continuous Titration Coordinates. Proteins 56:738–752.

43. Khandogin, J., and C. L. Brooks Iii. 2005. Constant pH Molecular Dynamics with Proton Tautomerism. Biophys. J. 89:141–157.

44. Wallace, J. A., and J. K. Shen. 2011. Continuous Constant pH Molecular Dynamics in Explicit Solvent with pH-Based Replica Exchange. J. Chem. Theory Comput. 7:2617–2629.

45. Chen, W., Y. Huang, and J. Shen. 2016. Conformational Activation of a Transmembrane Proton Channel from Constant pH Molecular Dynamics. J. Phys. Chem. Lett. 7:3961–3966.

46. Chen, W., B. H. Morrow, C. Y. Shi, and J. K. Shen. 2014. Recent development and application of constant pH molecular dynamics. Mol. Simul. 40:830–838.

47. Torrie, G. M., and J. P. Valleau. 1977. Nonphysical Sampling Distributions in Monte Carlo Free-Energy Estimation: Umbrella Sampling. J. Comput. Phys. 23:187–199.

48. Sugita, Y., A. Kitao, and Y. Okamoto. 2000. Multidimensional replica-exchange method for free-energy calculations. J. Chem. Phys. 113:6042–6051.

49. Zhang, C., D. Bell, M. Harger, and P. Ren. 2017. Polarizable Multipole-Based Force Field for Aromatic Molecules and Nucleobases. J. Chem. Theory Comput. 13:666–678.

50. Rupakheti, C. R., B. Roux, F. Dehez, and C. Chipot. 2018. Modeling induction phenomena in amino acid cation-π interactions. Theor. Chem. Acc. 137:174.

51. Inakollu, V. S. S., D. P. Geerke, C. N. Rowley, and H. Yu. 2020. Polarisable force fields: what do they add in biomolecular simulations? Curr. Opin. Struct. Biol. 61:182–190.

52. Klauda, J. B., R. M. Venable, J. A. Freites, J. W. O’Connor, D. J. Tobias, C. Mondragon-Ramirez, I. V. Vorobyov, A. D. MacKerell Jr., and R. W. Pastor. 2010. Update of the CHARMM All-Atom Additive Force Field for Lipids: Validation on Six Lipid Types. J. Phys. Chem. B 114:7830–7843.

53. Best, R. B., X. Zhu, J. Shim, P. E. M. Lopes, J. Mittal, M. Feig, and A. D. MacKerell Jr. 2012. Optimization of the Additive CHARMM All-Atom Protein Force Field Targeting Improved Sampling of the Backbone f, ? and Side-Chain χ1 and χ2 Dihedral Angles. J. Chem. Theory Comput. 8:3257–3273.

54. Huang, J., S. Rauscher, G. Nawrocki, T. Ran, M. Feig, B. L. de Groot, H. Grubmüller, and A. D. MacKerell Jr. 2017. CHARMM36m: an improved force field for folded and intrinsically disordered proteins. Nat. Methods 14:71–73.

55. Lamoureux, G., E. Harder, I. V. Vorobyov, B. Roux, and A. D. MacKerell Jr. 2006. A polarizable model of water for molecular dynamics simulations of biomolecules. Chem. Phys. Lett. 418:245–249.

56. Lamoureux, G., and B. Roux. 2006. Absolute Hydration Free Energy Scale for Alkali and Halide Ions Established from Simulations with a Polarizable Force Field. J. Phys. Chem. B 110:3308–3322.

57. Yu, H., T. W. Whitfield, E. Harder, G. Lamoureux, I. V. Vorobyov, V. M. Anisimov, A. D. MacKerell Jr., and B. Roux. 2010. Simulating Monovalent and Divalent Ions in Aqueous Solution Using a Drude Polarizable Force Field. J. Chem. Theory Comput. 6:774–786.

58. Chowdhary, J., E. Harder, P. E. M. Lopes, L. Huang, A. D. MacKerell Jr., and B. Roux. 2013. A Polarizable Force Field of Dipalmitoylphosphatidylcholine Based on the Classical Drude Model for Molecular Dynamics Simulations of Lipids. J. Phys. Chem. B 117:9142–9160.

59. Lopes, P. E. M., J. Huang, J. Shim, Y. Luo, H. Li, B. Roux, and A. D. MacKerell Jr. 2013. Polarizable Force Field for Peptides and Proteins Based on the Classical Drude Oscillator. J. Chem. Theory Comput. 9:5430– 5449.

60. Li, H., J. Chowdhary, L. Huang, X. He, A. D. MacKerell Jr., and B. Roux. 2017. Drude Polarizable Force Field for Molecular Dynamics Simulations of Saturated and Unsaturated Zwitterionic Lipids. J. Chem. Theory Comput. 13:4535–4552.

61. Lin, F.-Y., and A. D. MacKerell Jr. 2020. Improved Modeling of Cation-π and Anion-Ring Interactions Using the Drude Polarizable Empirical Force Field for Proteins. J. Comput. Chem. 41:439–448.

62. Yue, Z., W. Chen, H. I. Zgurskaya, and J. Shen. 2017. Constant pH Molecular Dynamics Reveals How Proton Release Drives the Conformational Transition of a Transmembrane Efflux Pump. J. Chem. Theory Comput. 13:6405–6414.

63. Yue, Z., C. Li, G. A. Voth, and J. M. J. Swanson. 2019. Dynamic Protonation Dramatically Affects the Membrane Permeability of Drug-like Molecules. J. Am. Chem. Soc. 141:13421–13433.

64. Durell, S. R., B. R. Brooks, and A. Ben-Naim. 1994. Solvent-Induced Forces between Two Hydrophilic Groups. J. Phys. Chem. 98:2198–2202.

65. Jorgensen, W. L., J. Chandrasekhar, J. D. Madura, R. W. Impey, and M. L. Klein. 1983. Comparison of simple potential functions for simulating liquid water. J. Chem. Phys. 79:926–935.

66. Senn, H. M., D. O’Hagan, and W. Thiel. 2005. Insight into Enzymatic C-F Bond Formation from QM and QM/MM Calculations. J. Am. Chem. Soc. 127:13643–13655.

67. Laage, D., H. Demirdjian, and J. T. Hynes. 2005. Intermolecular vibration-vibration energy transfer in solution: Hydrogen fluoride in water. Chem. Phys. Lett. 405:453–458.

68. MacKerell Jr., A. D., D. Bashford, M. Bellott, R. L. Dunbrack Jr., J. D. Evanseck, M. J. Field, S. Fischer, J. Gao, H. Guo, S. Ha, D. Joseph-McCarthy, L. Kuchnir, K. Kuczera, F. T. Lau, C. Mattos, S. Michnick, T. Ngo, D. T. Nguyen, B. Prodhom, W. E. Reiher, B. Roux, M. Schlenkrich, J. C. Smith, R. Stote, J. Straub, M. Watanabe, J. Wiórkiewicz-Kuczera, D. Yin, and M. Karplus. 1998. All-Atom Empirical Potential for Molecular Modeling and Dynamics Studies of Proteins. J. Phys. Chem. B 102:3586–3616.

69. Mackerell Jr., A. D., M. Feig, and C. L. Brooks Iii. 2004. Extending the Treatment of Backbone Energetics in Protein Force Fields: Limitations of Gas-Phase Quantum Mechanics in Reproducing Protein Conformational Distributions in Molecular Dynamics Simulations. J. Comput. Chem. 25:1400–1415.

70. Jo, S., T. Kim, and W. Im. 2007. Automated Builder and Database of Protein/Membrane Complexes for Molecular Dynamics Simulations. PLoS ONE 2:e880.

71. Jo, S., T. Kim, V. G. Iyer, and W. Im. 2008. CHARMM-GUI: A Web-Based Graphical User Interface for CHARMM. J. Comput. Chem. 29:1859–1865.

72. Jo, S., J. B. Lim, J. B. Klauda, and W. Im. 2009. CHARMM-GUI Membrane Builder for Mixed Bilayers and Its Application to Yeast Membranes. Biophys. J. 97:50–58.

73. Wu, E. L., X. Cheng, S. Jo, H. Rui, K. C. Song, E. M. Dávila-Contreras, Y. Qi, J. Lee, V. Monje-Galvan, R. M. Venable, J. B. Klauda, and W. Im. 2014. CHARMM-GUI Membrane Builder Toward Realistic Biological Membrane Simulations. J. Comput. Chem. 35:1997–2004.

74. Lee, J., X. Cheng, J. M. Swails, M. S. Yeom, P. K. Eastman, J. A. Lemkul, S. Wei, J. Buckner, J. C. Jeong, Y. Qi, S. Jo, V. S. Pande, D. A. Case, C. L. Brooks III, A. D. MacKerell Jr., J. B. Klauda, and W. Im. 2016. CHARMM-GUI Input Generator for NAMD, GROMACS, AMBER, OpenMM, and CHARMM/OpenMM Simulations Using the CHARMM36 Additive Force Field. J. Chem. Theory Comput. 12:405–413.

75. Abraham, M. J., T. Murtola, R. Schulz, S. Páll, J. C. Smith, B. Hess, and E. Lindahl. 2015. GROMACS: High performance molecular simulations through multi-level parallelism from laptops to supercomputers. SoftwareX 1–2:19–25.

76. Brooks, B. R., C. L. Brooks Iii, A. D. Mackerell, L. Nilsson, R. J. Petrella, B. Roux, Y. Won, G. Archontis, C. Bartels, S. Boresch, A. Caflisch, L. Caves, Q. Cui, A. R. Dinner, M. Feig, S. Fischer, J. Gao, M. Hodoscek, W. Im, K. Kuczera, T. Lazaridis, J. Ma, V. Ovchinnikov, E. Paci, R. W. Pastor, C. B. Post, J. Z. Pu, M. Schaefer, B. Tidor, R. M. Venable, H. L. Woodcock, X. Wu, W. Yang, D. M. York, and M. Karplus. 2009. CHARMM: The Biomolecular Simulation Program. J. Comput. Chem. 30:1545–1614.

77. Phillips, J. C., R. Braun, W. Wang, J. Gumbart, E. Tajkhorshid, E. Villa, C. Chipot, R. D. Skeel, L. Kalé, and K. Schulten. 2005. Scalable Molecular Dynamics with NAMD. J. Comput. Chem. 26:1781–1802.

78. Thurlkill, R. L., G. R. Grimsley, J. M. Scholtz, and C. N. Pace. 2006. pK values of the ionizable groups of proteins. Protein Sci. 15:1214–1218.

79. Lin, F.-Y., and A. D. MacKerell Jr. 2019. Improved Modeling of Halogenated Ligand–Protein Interactions Using the Drude Polarizable and CHARMM Additive Empirical Force Fields. J. Chem. Inf. Model. 59:215– 228.

80. Woolf, T. B., and B. Roux. 1994. Conformational Flexibility of o-Phosphorylcholine and o-Phosphorylethanolamine: A Molecular Dynamics Study of Solvation Effects. J. Am. Chem. Soc. 116:5916– 5926.

81. Vanýsek, P. 2006. Ionic Conductivity and Diffusion at Infinite Dilution. In CRC Handbook of Chemistry and Physics, 87th ed.; R. L. David, Ed. CRC Press, Boca Raton, FL; pp. 76–78.

82. Wang, J. H. 1954. Effect of Ions on the Self-Diffusion and Structure of Water in Aqueous Electrolytic Solutions. J. Phys. Chem. 58:686–692.

83. Prajapati, J. D., C. Mele, M. A. Aksoyoglu, M. Winterhalter, and U. Kleinekathöfer. 2020. Computational Modeling of Ion Transport in Bulk and through a Nanopore Using the Drude Polarizable Force Field. J. Chem. Inf. Model. 60:3188–3203.

84. Szabo, A., K. Schulten, and Z. Schulten. 1980. First passage time approach to diffusion controlled reactions. J. Chem. Phys. 72:4350–4357.

85. Hodgkin, A. L., and R. D. Keynes. 1955. The potassium permeability of a giant nerve fibre. J. Physiol. 128:61– 88.

86. Bernèche, S., and B. Roux. 2001. Energetics of ion conduction through the K+ channel. Nature 414:73–77.

87. Bernèche, S., and B. Roux. 2003. A microscopic view of ion conduction through the K+ channel. Proc. Natl. Acad. Sci. U.S.A. 100:8644–8648.

88. Egwolf, B., and B. Roux. 2010. Ion Selectivity of the KcsA Channel: A Perspective from Multi-Ion Free Energy Landscapes. J. Mol. Biol. 401:831–842.

89. Medovoy, D., E. Perozo, and B. Roux. 2016. Multi-ion free energy landscapes underscore the microscopic mechanism of ion selectivity in the KcsA channel. Biochim. Biophys. Acta 1858:1722–1732.

90. Muth, T. R., and S. Schuldiner. 2000. A membrane-embedded glutamate is required for ligand binding to the multidrug transporter EmrE. EMBO J. 19:234–240.

91. Yerushalmi, H., and S. Schuldiner. 2000. An Essential Glutamyl Residue in EmrE, a Multidrug Antiporter from Escherichia coli. J. Biol. Chem. 275:5264–5269.

92. Emsley, J. 1980. Very Strong Hydrogen Bonding. Chem. Soc. Rev. 9:91–124.

93. Chowdhuri, S., and A. Chandra. 2006. Dynamics of Halide Ion-Water Hydrogen Bonds in Aqueous Solutions: Dependence on Ion Size and Temperature. J. Phys. Chem. B 110:9674–9680.

94. Allen, T. W., O. S. Andersen, and B. Roux. 2004. Energetics of ion conduction through the gramicidin channel. Proc. Natl. Acad. Sci. U.S.A. 101:117–122.

95. Allen, T. W., O. S. Andersen, and B. Roux. 2006. Molecular dynamics — potential of mean force calculations as a tool for understanding ion permeation and selectivity in narrow channels. Biophys. Chem. 124:251–267.

96. Peng, Y., J. M. J. Swanson, S. Kang, R. Zhou, and G. A. Voth. 2015. Hydrated Excess Protons Can Create Their Own Water Wires. J. Phys. Chem. B 119:9212–9218.

97. Li, C., and G. A. Voth. 2021. A quantitative paradigm for water-assisted proton transport through proteins and other confined spaces. Proc. Natl. Acad. Sci. U.S.A. 118:e2113141118.

98. Ariz-Extreme, I., and J. S. Hub. 2018. Assigning crystallographic electron densities with free energy calculations-The case of the fluoride channel Fluc. PLoS ONE 13:e0196751.

99. Jackson, M. R., R. Beahm, S. Duvvuru, C. Narasimhan, J. Wu, H. N. Wang, V. M. Philip, R. J. Hinde, and E. E. Howell. 2007. A Preference for Edgewise Interactions between Aromatic Rings and Carboxylate Anions: The Biological Relevance of Anion-Quadrupole Interactions. J. Phys. Chem. B 111:8242–8249.

100. Nosé, S. 1984. A molecular dynamics method for simulations in the canonical ensemble. Mol. Phys. 52:255– 268.

101. Hoover, W. G. 1985. Canonical dynamics: Equilibrium phase-space distributions. Phys. Rev. A 31:1695–1697.

102. Parrinello, M., and A. Rahman. 1981. Polymorphic transitions in single crystals: A new molecular dynamics method. J. Appl. Phys. 52:7182–7190.

103. Olsson, M. H. M., C. R. Søndergaard, M. Rostkowski, and J. H. Jensen. 2011. PROPKA3: Consistent Treatment of Internal and Surface Residues in Empirical pKa Predictions. J. Chem. Theory Comput. 7:525– 537.

104. Søndergaard, C. R., M. H. M. Olsson, M. Rostkowski, and J. H. Jensen. 2011. Improved Treatment of Ligands and Coupling Effects in Empirical Calculation and Rationalization of pKa Values. J. Chem. Theory Comput. 7:2284–2295.

105. Hess, B., H. Bekker, H. J. C. Berendsen, and J. G. E. M. Fraaije. 1997. LINCS: A Linear Constraint Solver for Molecular Simulations. J. Comput. Chem. 18:1463–1472.

106. Darden, T., D. York, and L. Pedersen. 1993. Particle mesh Ewald: An N•log(N) method for Ewald sums in large systems. J. Chem. Phys. 98:10089–10092.

107. Essmann, U., L. Perera, M. L. Berkowitz, T. Darden, H. Lee, and L. G. Pedersen. 1995. A smooth particle mesh Ewald method. J. Chem. Phys. 103:8577–8593.

108. Wang, Z., J. M. J. Swanson, and G. A. Voth. 2018. Modulating the Chemical Transport Properties of a Transmembrane Antiporter via Alternative Anion Flux. J. Am. Chem. Soc. 140:16535–16543.

109. Wang, Z., J. M. J. Swanson, and G. A. Voth. 2020. Local Conformational Dynamics Regulating Transport Properties of a Cl-/H+ Antiporter. J. Comput. Chem. 41:513–519.

110. Ryckaert, J.-P., G. Ciccotti, and H. J. C. Berendsen. 1977. Numerical Integration of the Cartesian Equations of Motion of a System with Constraints: Molecular Dynamics of n-Alkanes. J. Comput. Phys. 23:327–341.

111. Feller, S. E., Y. Zhang, R. W. Pastor, and B. R. Brooks. 1995. Constant pressure molecular dynamics simulation: The Langevin piston method. J. Chem. Phys. 103:4613–4621.

112. Im, W., M. S. Lee, and C. L. Brooks Iii. 2003. Generalized Born Model with a Simple Smoothing Function. J. Comput. Chem. 24:1691–1702.

113. Im, W., M. Feig, and C. L. Brooks III. 2003. An Implicit Membrane Generalized Born Theory for the Study of Structure, Stability, and Interactions of Membrane Proteins. Biophys. J. 85:2900–2918.

114. Chen, J., W. Im, and C. L. Brooks III. 2006. Balancing Solvation and Intramolecular Interactions: Toward a Consistent Generalized Born Force Field. J. Am. Chem. Soc. 128:3728–3736.

115. Miyamoto, S., and P. A. Kollman. 1992. SETTLE: An Analytical Version of the SHAKE and RATTLE Algorithm for Rigid Water Models. J. Comput. Chem. 13:952–962.

116. Liang, R., H. Li, J. M. J. Swanson, and G. A. Voth. 2014. Multiscale simulation reveals a multifaceted mechanism of proton permeation through the influenza A M2 proton channel. Proc. Natl. Acad. Sci. U.S.A. 111:9396–9401.

117. Hill, A. V. 1910. The possible effects of the aggregation of the molecules of haemoglobin on its dissociation curves. J. Physiol. 40:iv–vii.

118. Kumar, S., J. M. Rosenberg, D. Bouzida, R. H. Swendsen, and P. A. Kollman. 1992. The Weighted Histogram Analysis Method for Free-Energy Calculations on Biomolecules. I. The Method. J. Comput. Chem. 13:1011– 1021.

119. De Loof, H., L. Nilsson, and R. Rigler. 1992. Molecular Dynamics Simulation of Galanin in Aqueous and Nonaqueous Solution. J. Am. Chem. Soc. 114:4028–4035.

120. Kumar, S., and R. Nussinov. 2002. Close-Range Electrostatic Interactions in Proteins. ChemBioChem 3:604– 617.

121. Chakravarty, S., A. R. Ung, B. Moore, J. Shore, and M. Alshamrani. 2018. A Comprehensive Analysis of Anion-Quadrupole Interactions in Protein Structures. Biochemistry 57:1852–1867.

122. Humphrey, W., A. Dalke, and K. Schulten. 1996. VMD: Visual Molecular Dynamics. J. Mol. Graph. 14:33– 38.

