## Supplemental File for "Ion Permeation, Selectivity, and Electronic Polarization in Fluoride Channels"

### 1 Simulation Protocols

#### 1.1 System preparation

Generated using the CHARMM-GUI *Membrane Builder* (1-5), the simulation system consisted of a Fluc protein from *E. coli* (Fluc-Ec2, PDB: 5A43) (6), 135 1-palmitoyl-2-oleoyl-*sn*-glycero-3-phosphoethanolamine (POPE) lipids, 17 $K^+$ anions, 25 $Cl^-$ anions, and ~7,000 TIP3P water molecules in a 71 Å × 71 Å × 82 Å box. Because the force field parameters for $F^-$ are not included in the distributed additive CHARMM force field (7-11), the two crystal $F^-$ ions were replaced with $Cl^-$.

#### 1.2 System equilibration

The system was equilibrated using GROMACS (12) (version 2019.3) for 100 ns at a constant temperature of 310 K using the Nosé-Hoover thermostat (13,14) and a constant pressure of 1 atm using the Parrinello-Rahman barostat (15). The CHARMM36m (10,11), CHARMM36 (9), and CHARMM-modified (16) TIP3P (17) models were used to represent protein, lipids, and waters, respectively. Based on p$K_a$s predicted by PROPKA3.1 (18,19), both E86s were protonated, while all other residues were in their standard protonation states. The leapfrog integrator was used to propagate the simulation. LINCS (20) was used to restrain bonds involving hydrogen atoms to allow for a 2-fs step length. The particle-mesh Ewald (PME) method (21,22) with a real-space cutoff of 12.0 Å was applied to calculate electrostatic interactions. A switching function was applied to the van der Waals (vdW) force from 10.0 to 12.0 Å. The neighbor list was updated every 20 MD steps. All heavy atoms of the protein and two bound ions were restrained using a harmonic potential of 1.00 kcal/(mol•Å$^2$). Other simulation details are consistent with our previous work (23,24).

#### 1.3 Continuous constant-pH molecular dynamics (CpHMD) simulations

The last snapshot from the GROMACS equilibration was retrieved as the starting configuration, with all $Cl^-$ ions back-converted to $F^-$ and the crystallographic $Na^+$ docked into the protein. Simulations were conducted using CHARMM (25) (version c42b2) and utilized the CHARMM27/CMAP (7,8) force field to model protein. The parameters for $F^-$ (Appendix I) and HF (Appendix II) were reported by Senn *et al.* (26) and Laage *et al.* (27), respectively. The CpHMD parameters for HF (Appendix III and **Fig. S10**) were derived using the protocol described by Lee *et al.* (28) and detailed in Yue *et al.* (29). The leapfrog Verlet integrator was used to propagate the spatial coordinates. The vdW potential was switched from 10 to 12 Å in aqueous CpHMD simulations, and the vdW force was switched from 8 to 12 Å in membrane simulations. Electrostatic interactions were computed using PME (21,22) with a real-space cutoff of 12 Å and a sixth-order interpolation with 1-Å grid spacing. SHAKE (30) was used to restrain bonds involving hydrogen atoms to allow for a 2-fs step length. All simulations were conducted with periodic boundary conditions at a constant temperature of 310 K by the Nosé-Hoover thermostat (13,14) and a constant pressure of 1 atm by the Langevin piston pressure-coupling algorithm (31).

In the membrane-enabled (32) hybrid-solvent (33) CpHMD simulations (28,34), the conformational dynamics was propagated using explicit solvent and lipid molecules, while the electrostatic hydration forces on the titration coordinates were computed using implicit solvent and membrane described by the generalized-Born (GB) model GBSW (35,36) with optimized GB input radii by Chen *et al.* (37). The implicit membrane was represented by an infinite low-dielectric-constant slab harboring a high-dielectric-constant exclusion cylinder that aligns with the Z-axis to mimic the protein region (36). The implicit-membrane thickness was 35 Å, and the radius of the exclusion cylinder was 15 Å. The dielectric constant was switched from 2 to 80 within 2.5 Å for both leaflets. The ionic strength in the GB calculation was set to 0.150 M. Following Wallace *et al.* (33), the fictitious lambda particles with a mass of 10 atomic mass units were propagated using the Langevin algorithm with a collision frequency of 5 $ps^{-1}$, the titration coordinates were updated every 10 MD steps to allow for water relaxation. All Asp, Glu, His, Arg19 residues, and two HF ligands were allowed to titrate. A spherical restraint was applied via the MMFP utility in CHARMM (25) to the center-of-mass (COM) of the protein heavy atoms with a force constant of 50 kcal/(mol•$Å^2$). Ions were excluded from the hydrophobic membrane region (–17.5 Å < Z < 17.5 Å) by a planar restraint of 1 kcal/(mol•$Å^2$). The HF in chain B was restrained around its initial location by a spherical restraint of 2 kcal/(mol•$Å^2$). In the CpHMD equilibrations at pH 7, a planar restraint of 10 kcal/(mol•$Å^2$) was applied to the Z-component of the COM of the HF in chain A (ZCOM) to relocate it to the designated location along the membrane normal. Twenty-one replicates of the starting configuration were thus generated by constraining the ZCOM from –15 Å to 15 Å with a 1.5 Å interval. The CpHMD equilibrations ran for 50 ns until the backbone RMSD of the protein plateaued (**Fig. S11**). When the pH-based replica exchange (pH-REX) (33) was applied, exchanges between neighboring replicas were attempted every 500 MD steps.

#### 1.4 Replica-exchange umbrella sampling (REUS) simulations

The final snapshots of the 21 replicates from the CpHMD equilibrations were retrieved and converted to fixed-charge-state style by assigning the protonation states of the ionizable residues at pH 7 based on the $pK_a$s predicted from the pH-REX CpHMD simulation (**Table 1**) and by adjusting the number of cations to make the systems charge neutral. The replica-exchange umbrella sampling (REUS) (38) simulations were then performed to calculate the potentials of mean force (PMFs) of $F^-$ permeation, using the NAMD simulation package (39).

In simulations using polarizable force field, the CHARMM Drude-2013 polarizable force field (40-45) was employed to model the protein (44), lipids (43,45), ions (41,42), and SWM4-NDP waters (40). The cation-π correction (46) was added to better describe the interactions between protonated His106 and Phe83. In nonpolarizable simulations, the CHARMM36m (9-11) force field was used to model the protein and lipids, and the CHARMM-modified (16) TIP3P (17) water model was adopted. The step length was 0.25 and 2 fs for polarizable and nonpolarizable simulations, respectively. The PME method (21) was used to treat the long-range electrostatics, with a real-space cutoff of 12.0 Å and an accuracy threshold of $10^{-6}$. A switching function was applied to vdW and the short-range electrostatics from 10 to 12 Å. The neighbor list was updated every 10 MD steps using a cutoff distance of 16.0 Å. The SETTLE algorithm (47) was used to restrain bonds involving hydrogen atoms. In aqueous bulk, cylindrical restraints were applied to restrain the $F^-$ within the same region on the XY-plane as within the protein, following our previous work (48). Other simulation settings were consistent with our previous work (23,24). The collective variable (CV) was the Z-component of the vector from the COM of α-carbon (Cα) atoms of the protein to the $F^-$ ion.

### 2 Analysis Protocols

#### 2.1 Calculation of p$K_a$

The p$K_a$ was computed by fitting the unprotonated fraction $S^{\mathrm{unprot}}$ *vs.* pH to the Hill equation (49),

$$S^{\mathrm{unprot}} = \frac{1}{1+10^{n(\mathrm{p}K_a-\mathrm{pH})}}, \quad \text{(S1)}$$

where the Hill coefficient *n* describes the steepness of the transition region in a titration curve. $S^{\mathrm{unprot}}$ was determined by counting the population of protonated (defined as those with $\lambda^{\mathrm{HF}} \leq 0.2$) and deprotonated ($\lambda^{\mathrm{HF}} \geq 0.8$) states for every pH.

#### 2.2 Calculation of fluoride permeation free energy

The Weighted Histogram Analysis Method (WHAM) (50) was utilized to construct the potential of mean force (PMF). The error of PMF was estimated as the standard deviation from block analysis.

#### 2.3 Calculation of fluoride conductance

The effective rate constant $k$ for the $F^-$ permeation was calculated from the PMF $F(z)$ based on the mean first passage time (51):

$$k = \left\{ \int_{z_a}^{z_b} dz' D(z')^{-1} \exp[F(z')/k_B T] \int_{z_a}^{z'} dz'' \exp[-F(z'')/k_B T] \right\}^{-1}, \quad \text{(S2)}$$

where $z_a = -20$ Å and $z_b = 20$ Å are the entrance and the exit of the $F^-$ pore, $k_B$ is the Boltzmann constant, $T$ is the temperature, $D(z)$ is the local diffusion constant given by the Woolf-Roux equation (52):

$$D(z_i) = \lim_{s\to 0} \frac{-\hat{C}(s;z_i)\langle \delta z^2\rangle_{(i)}\langle \dot{z}^2\rangle_{(i)}}{\hat{C}(s;z_i)\left[s\langle \delta z^2\rangle_{(i)} + \langle \dot{z}^2\rangle_{(i)}/s\right] - \langle \delta z^2\rangle_{(i)}\langle \dot{z}^2\rangle_{(i)}}, \quad \text{(S3)}$$

where $\delta z = z - z_i$ is the deviation from the center, $C(t;z_i) = \langle \dot{z}(t)\dot{z}(0)\rangle_i$ is the position-dependent velocity autocorrelation function for the $F^-$ permeation along the Z-axis, calculated from the *i*-th umbrella "window", $z_i$ is the reference point for the harmonic restraint potential in window $i$, and $\hat{C}(s;z_i) = \int_0^\infty e^{-st} C(t;z_i)dt$ is the Laplace transform of the autocorrelation function. To value at the limit of $s \to 0$ was linearly extrapolated from the range 0.2 (MD step)$^{-1} \leq s \leq 0.3$ (MD step)$^{-1}$. The errors of the aqueous *D* and the *k* were also estimated from block analysis and reported as standard deviations.

#### 2.4 Other analyses

All other analyses were conducted using CHARMM (version c42b2) (25). In general, COMs were calculated using the *coor stat* command. Atomic distances and angles were computed using the *quick* command. The minimal distances between two groups of atoms were computed using the *coor mind* command.

Hydration numbers ($N_{\mathrm{hydr}}$) were defined as the number of water oxygen atoms within 3.5 and 3.0 Å (first solvation shell, Refs. (41,42)) respectively for $F^-$ and $Na^+$ and computed using the *coor anal* command.

The coordination number ($N_{\mathrm{coord}}$) of the central $Na^+$, defined as the number of oxygen atoms within 3.0 Å, was computed using the *scalar* command.

The number of hydrogen-bonds between $F^-$ and Fluc-Ec2 ($N_{\mathrm{HBond}}$) was calculated using the *coor hbond* command based on the donor–H cutoff distance of 2.4 Å (53). When calculating the occupancy of a specific HBond, both the donor–acceptor cutoff distance of 3.5 Å and the donor–H–acceptor cutoff angle of 150° were applied.

Ion-pair (or salt-bridge) was considered to form if the minimum distance between the positively and negatively charged atoms is ≤ 4.0 Å (54).

Anion-π pair was present if $F^-$ is within 4.5 Å of any ring carbon atoms and subtends an angle of ≤ 35° with the ring plane (55). The angle was derived from the relative orientation between the ring normal (calculated using the *coor lsqp* command) and the vector connecting $F^-$ and the ring center (calculated using the *coor axis* command).

Following Lin *et al.* (56), the interaction energy between $F^-$ and Fluc ($E_{inter}$) was calculated using

$$E_{\mathrm{inter}} = E_{\mathrm{comp}} - E_{\mathrm{prot}} - E_{\mathrm{lig}}, \tag{S4}$$

where $E_{comp}$, $E_{prot}$, and $E_{lig}$ represent the total energy of the protein–ligand complex, protein, and ligand, respectively. Note that the ligand corresponds to the $F^-$ in monomer A while the protein includes Fluc, the $F^-$ in monomer B and the central $Na^+$. $E_{inter}$ can also be written as

$$E_{\mathrm{inter}} = E_{\mathrm{vdW}} + E_{\mathrm{elec}} + E_{\mathrm{self}}, \tag{S5}$$

where $E_{vdW}$, $E_{elec}$, and $E_{self}$ respectively represent the vdW, Coulombic and electronic self-polarization energy. Note that $E_{self}$ is only included in Drude. For C36m simulations, $E_{inter}$ was directly calculated from the raw trajectories using the *inter* command. For Drude simulations, relaxation of the electronic degree of freedom before energy evaluation is required. To do so, the Drude particles were minimized first by steepest descent with a step size of 0.01 to a force gradient of $10^{-2}$ kcal/(mol•Å) then by adopted basis Newton-Raphson with a step size of 0.02 to a force gradient of $10^{-5}$ kcal/(mol•Å), while fixing the real atoms. $E_{comp}$, $E_{prot}$, and $E_{lig}$ were then computed using the *inter* command. The trajectories excluding solvents were used.

The Drude particles and lone pairs were not included in all those analyses except the energy calculations.

#### 2.5 PDB survey

All Fluc crystal structures deposited in the PDB bank (**Table S1**) were collected. The PDB 5NKQ (6) was chosen as the reference structure and re-oriented with respect to the membrane using the PPM server (57). All other PDBs were aligned to 5NKQ by least-square fitting the Cα atoms. The membrane-positioned PDBs were then analyzed using the in-house Tcl scripts for VMD (58).

### 3 Supplementary Figures and Tables

#### 3.1 Figures and tables discussed in the main text

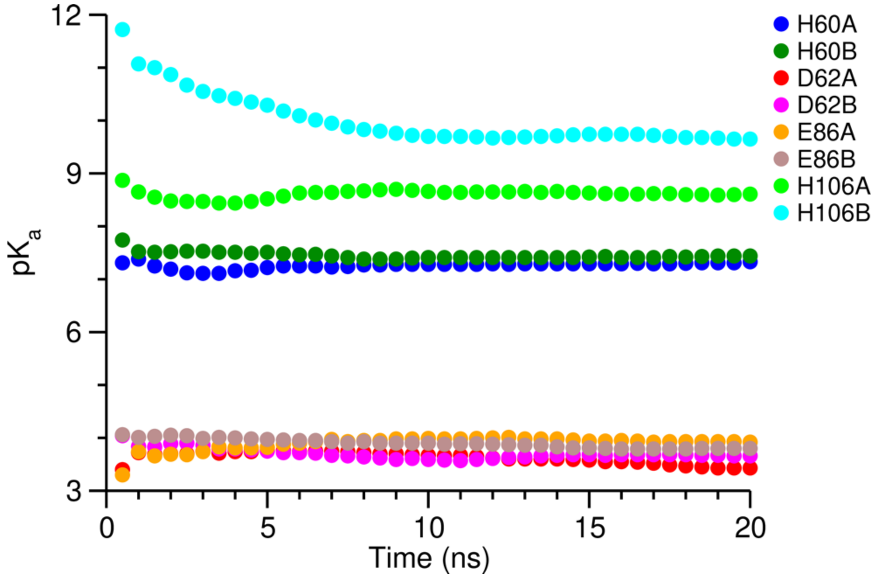


**Figure S1. Convergence of the p$K_a$ values in pH-REX CpHMD.** p$K_a$'s were calculated cumulatively versus simulation time. Note that the Fluc-bound fluorides and R19s stayed charged in the pH range investigated in the simulation (see **Table 1**).

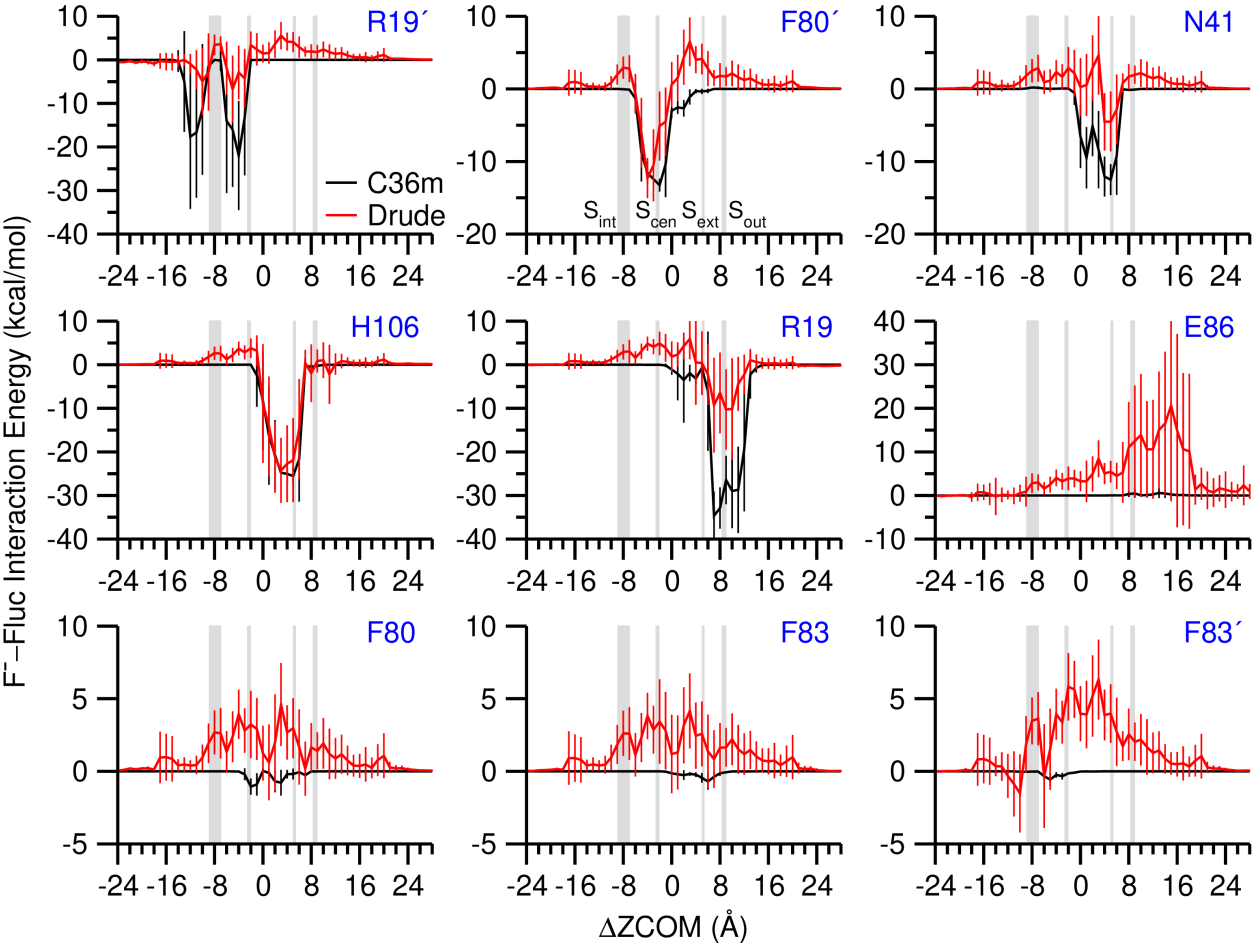


**Figure S2. Interaction energy of $F^-$ with essential Fluc-Ec2 residues.** The means and fluctuations (shown as error bars) were calculated every 1 Å along ΔZCOM. Grey boxes mark the crystallographic locations of $S_{int}$, $S_{cen}$, $S_{ext}$, and $S_{out}$ $F^-$.

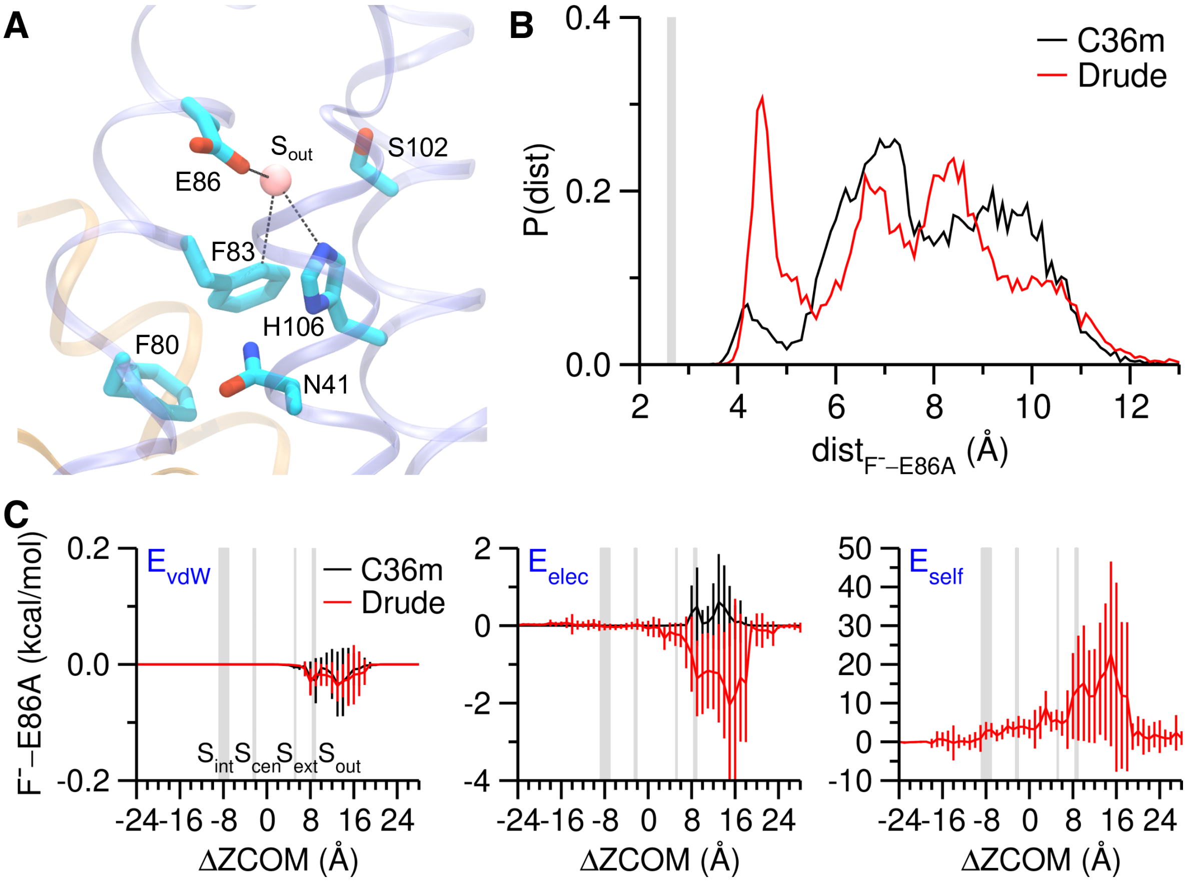


**Figure S3. Interaction between F⁻ and E86. (A)** $S_{out}$ F⁻ binding site with essential residues labeled (PDB: 6BX5). Black dashed lines indicate hydrogen-bond, ion-pair, and anion-π pair. **(B)** Distribution of the minimum distance between F⁻ and E86 side-chain oxygen atoms in states V and VI (ΔZCOM from 5 to 11 Å, see **Fig. 2A**). Grey box marks the value collected from PDB structures (centroid and halved box width respectively represent the mean and standard deviation). **(C)** Decomposition of the F⁻–E86 interaction energy. The means and fluctuations (shown as error bars) were calculated every 1 Å along ΔZCOM. Note $E_{self}$ is only included in Drude. Grey boxes mark the crystallographic locations of $S_{int}$, $S_{cen}$, $S_{ext}$, and $S_{out}$ F⁻.

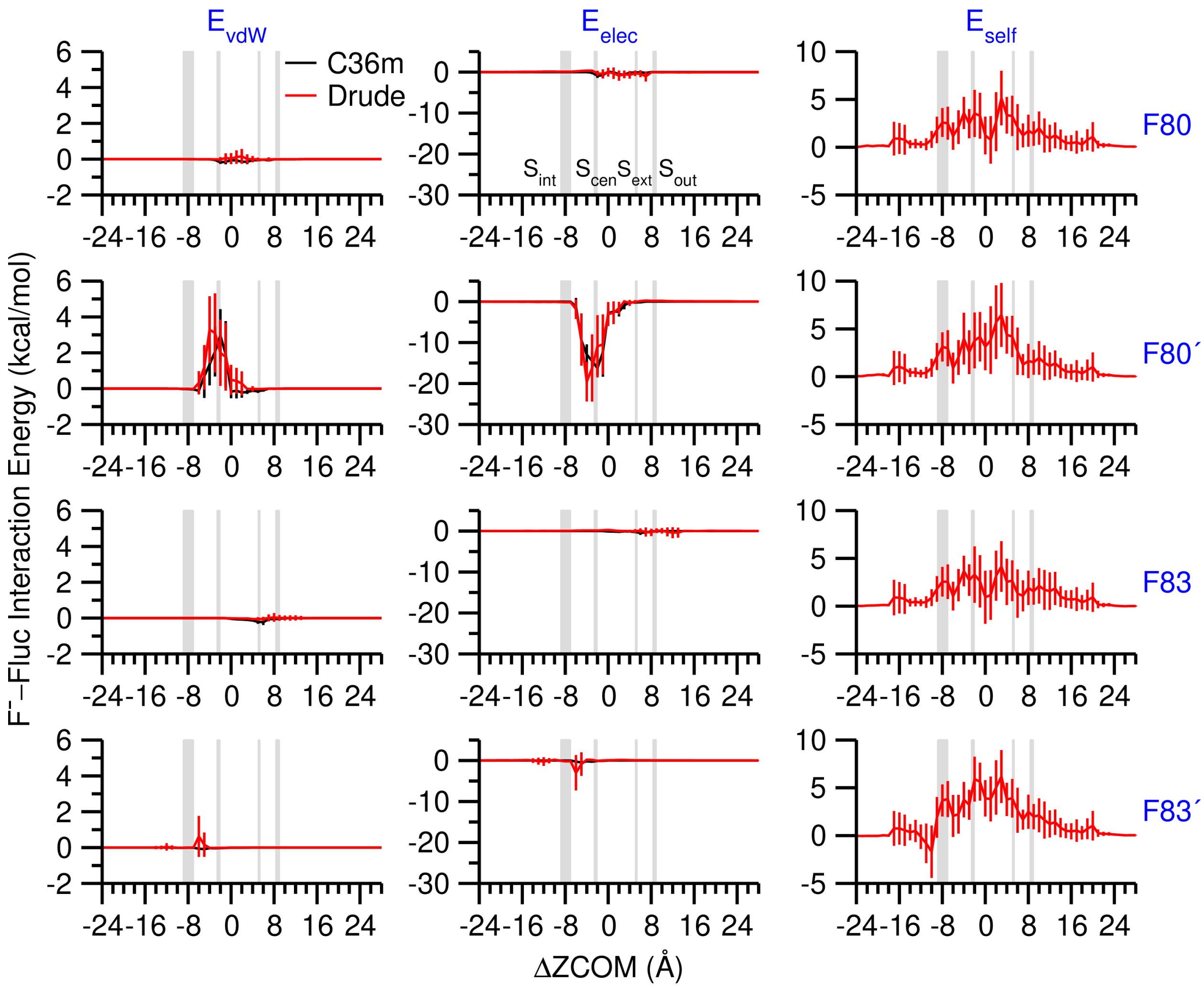


**Figure S4. Decomposition of the interaction energy between F⁻ and the "Phe-box".** The means and fluctuations (shown as error bars) were calculated every 1 Å along ΔZCOM. Note that $E_{self}$ is only included in Drude. Grey boxes mark the crystallographic locations of $S_{int}$, $S_{cen}$, $S_{ext}$, and $S_{out}$ F⁻.

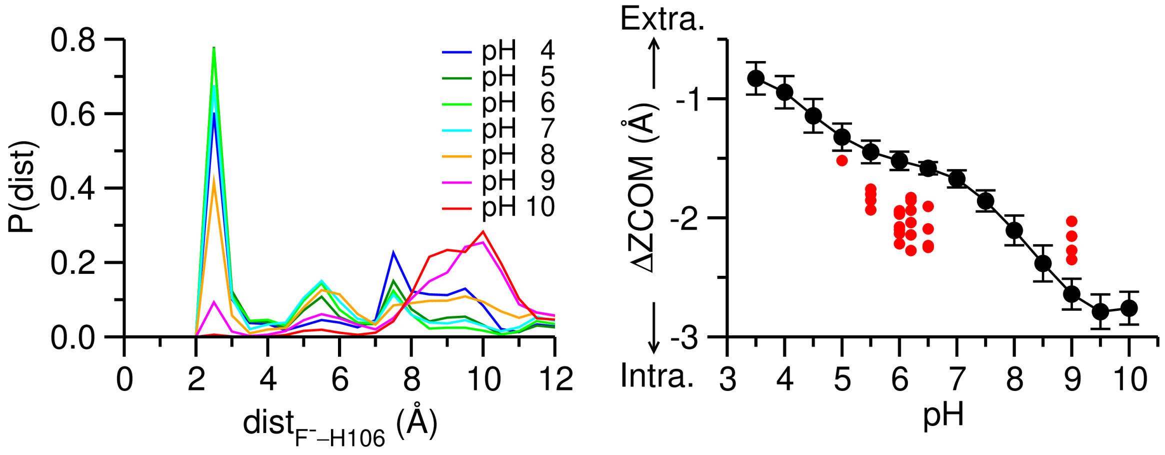


**Figure S5. Correlation between F⁻–H106 ion-pair and F⁻ location in pH-REX CpHMD. (Left)** Distribution of the minimal distance between F⁻ and H106 side-chain nitrogen atoms. The peak around 2–3 Å indicates the formation of a salt bridge (54). **(Right)** pH-dependent ΔZCOM of F⁻. Means and standard deviations (shown as error bars) were calculated from block analysis. Red circles indicate the values collected from crystal structures.

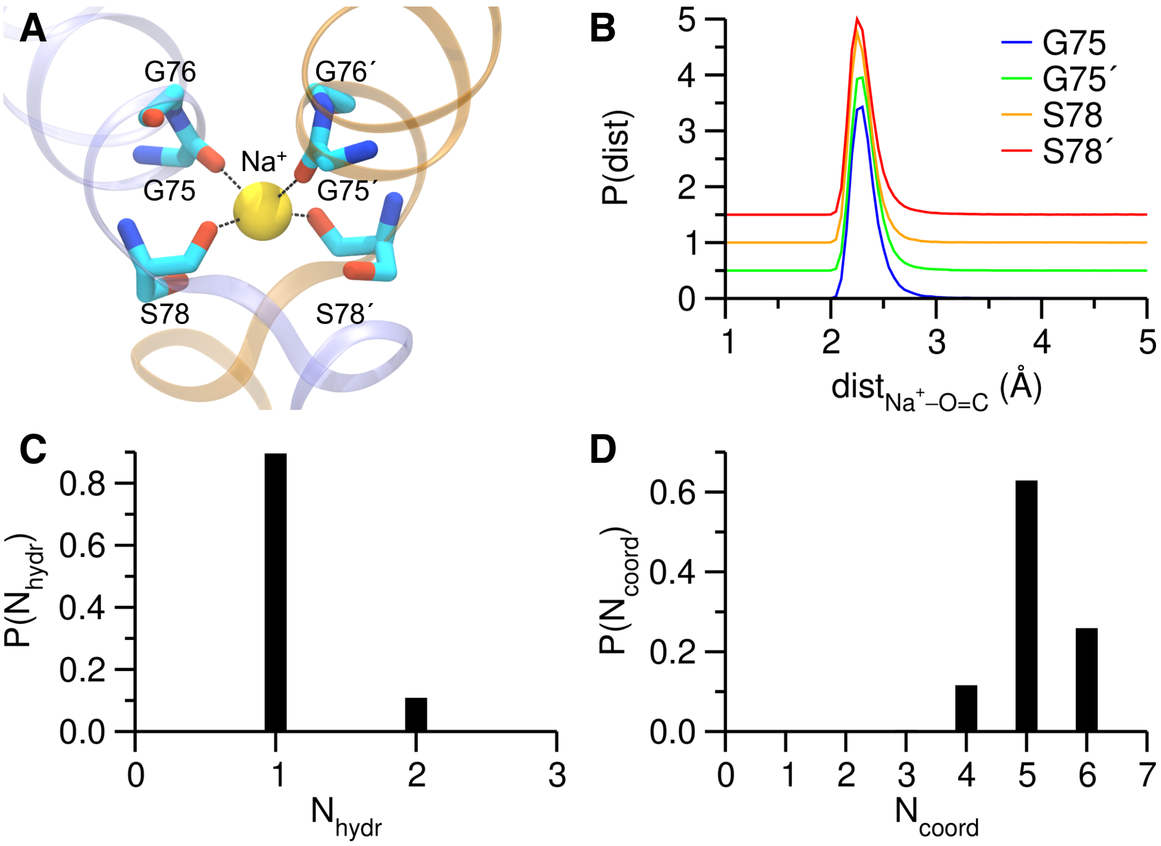


**Figure S6. Interaction between $Na^+$ and Fluc. (A)** $Na^+$ coordinated with the backbone carbonyl oxygen atoms of G75s and S78s (PDB: 5A43). Black dashed lines indicate interactions in the PDB structure. Note that our simulations found $Na^+$ could also interact with the backbone carbonyl oxygen of G76 and the side-chain hydroxyl oxygens of S78s. **(B)** Distribution of the distances between $Na^+$ and the backbone carbonyl oxygen atoms of G75s and S78s. The curves for G75´ and S78s were upshifted for clarity. The peak within 3 Å indicates coordination. Data was collected from all the "windows" in the Drude REUS simulations (same below). **(C)** Occupancy of $Na^+$ hydration numbers (the number of water oxygen atoms within 3.0 Å of $Na^+$). **(D)** Occupancy of $Na^+$ coordination numbers (the number of oxygen atoms within 3.0 Å of $Na^+$).

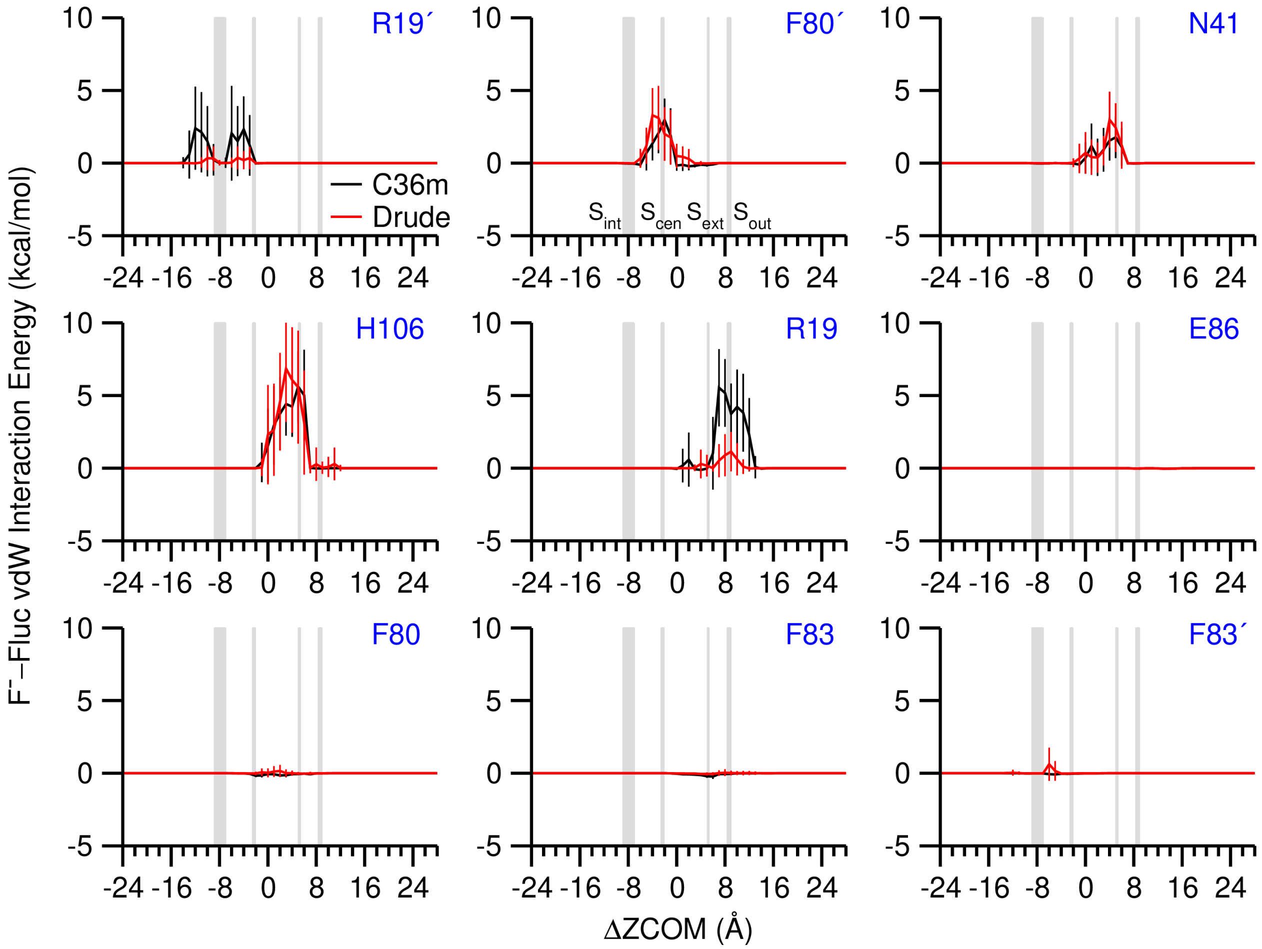


**Figure S7. The vdW interaction energy of F⁻ with essential Fluc-Ec2 residues.** The means and fluctuations (shown as error bars) were calculated every 1 Å along ΔZCOM. Grey boxes mark the crystallographic locations of $S_{int}$, $S_{cen}$, $S_{ext}$, and $S_{out}$ F⁻.

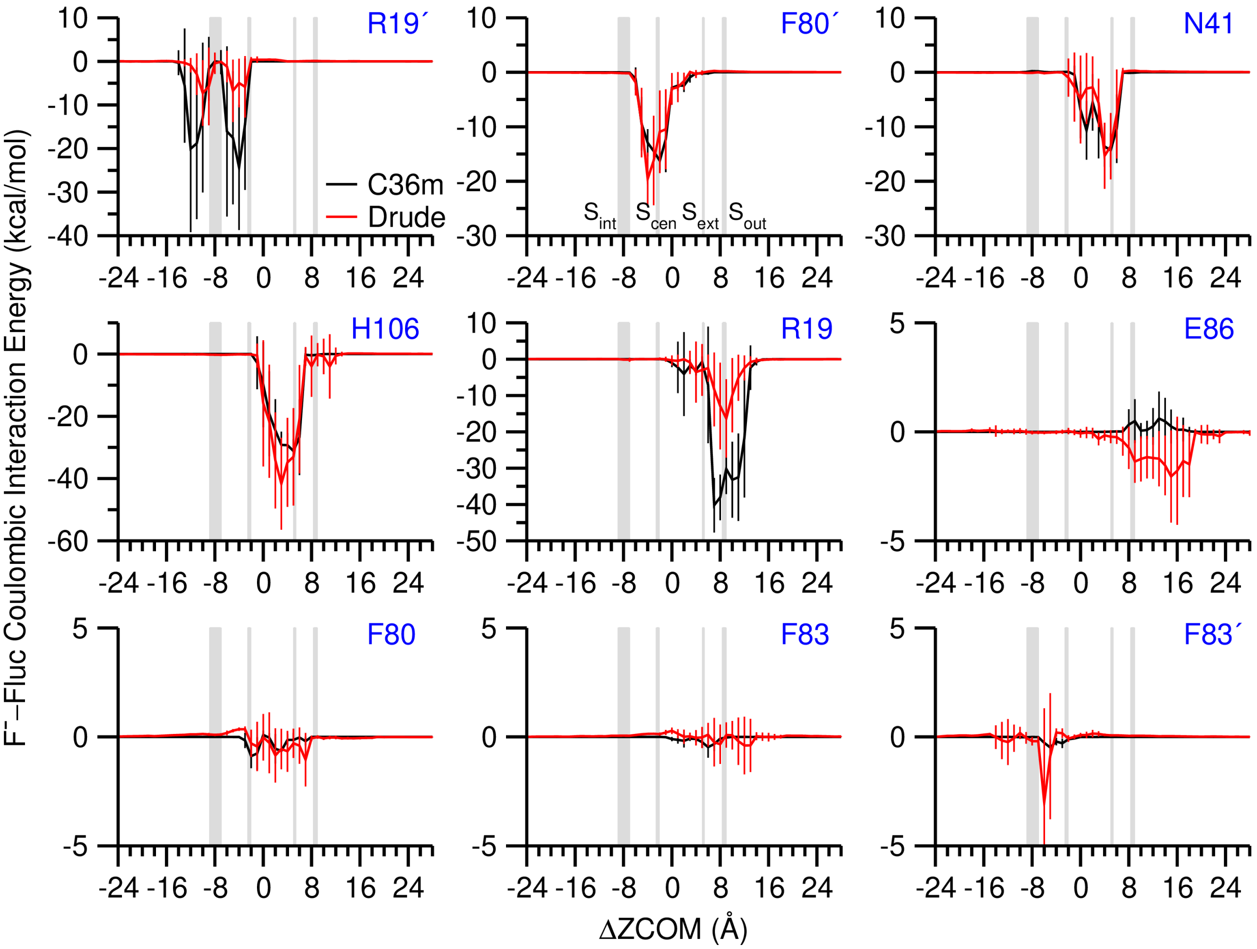


**Figure S8. The Coulombic interaction energy of F⁻ with essential Fluc-Ec2 residues.** The means and fluctuations (shown as error bars) were calculated every 1 Å along ΔZCOM. Grey boxes mark the crystallographic locations of $S_{int}$, $S_{cen}$, $S_{ext}$, and $S_{out}$ F⁻.

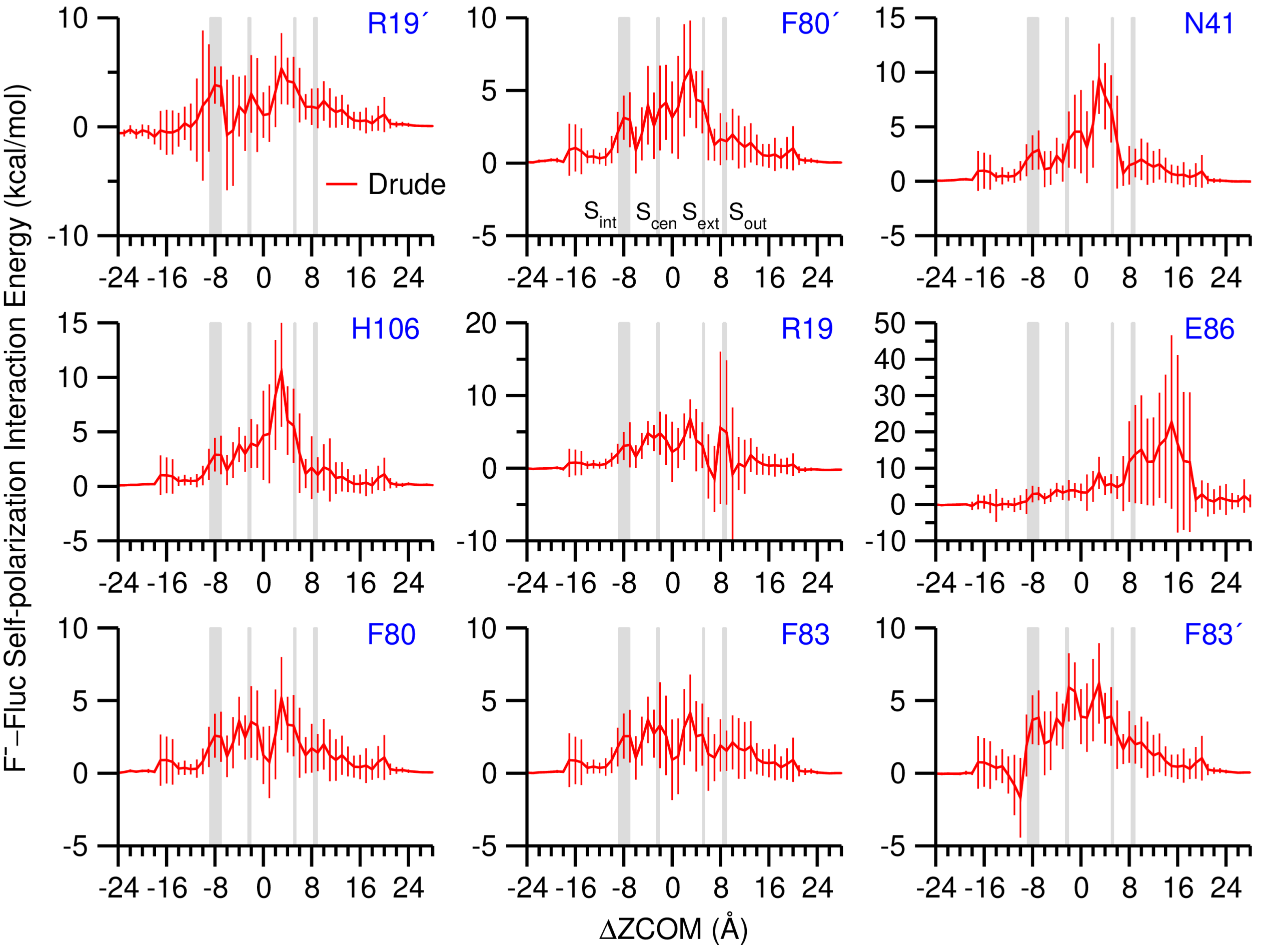


**Figure S9. The Drude electronic self-polarization interaction energy of $F^-$ with essential Fluc-Ec2 residues.** The means and fluctuations (shown as error bars) were calculated every 1 Å along ΔZCOM. Grey boxes mark the crystallographic locations of $S_{int}$, $S_{cen}$, $S_{ext}$, and $S_{out}$ $F^-$.

**Table S1. Comparison of published crystal structures of Fluc**

| PDB | Organism | Mutation[a] | Resolution (Å) | pH[b] | $F^-$ binding sites[c] | | | | Reference |
|---|---|---|---|---|---|---|---|---|---|
| | | | | | $S_{int}$ | $S_{cen}$ | $S_{ext}$ | $S_{out}$ | |
| 5A40 | B. pertusis | R29K/E94C | 3.60 | 8.5 | - | | | | Newstead (6) |
| 5NKQ | | | 2.17 | 5.5 | - | AB | AB | - | Newstead (6) |
| 6BQO | | | 2.80 | 5.0 | - | B | A | - | Stockbridge (59) |
| 5A43 | E. coli | R25K M1/MSE A51MSE M68/MSE M113/MSE | 2.58 | 6.0 | - | AB | - | - | Newstead (6) |
| 5KBN | | R25K/F80I | 2.48 | 6.2 | - | - | - | AB | Miller (60) |
| 5KOM | | R25K/F83I | 2.69 | 6.5 | - | AB | - | - | Miller (60) |
| 6B24 | | R25K/F80Y | 2.75 | 6.2 | - | AB | - | AB | Miller (61) |
| 6B2A | | R25K/F80M | 2.65 | 6.2 | - | AB | - | AB | Miller (61) |
| 6B2B | | R25K/F83M | 2.60 | 6.0 | - | AB | - | B | Miller (61) |
| 6B2D | | R25K/T114S | 3.01 | 6.5 | - | AB | - | B | Miller (61) |
| 6BX4 | | R25K | 2.55 | 6.0 | - | AB | - | AB | Miller (62) |
| 6BX5 | | R25K | 3.00 | 6.2 | - | AB | - | AB | Miller (62) |
| 7KK8[d] | | R25K/S81T | 2.70 | 6.0 | AB | B | - | - | Stockbridge (63) |
| 7KK9[d] | | R25K/S81A/T82A | 3.10 | 6.0 | AB | - | - | - | Stockbridge (63) |
| 7KKA[d] | | R25K/S81A | 2.50 | 9.0 | AB | AB | - | - | Stockbridge (63) |
| 7KKB[d] | | R25K/C74A/S81C | 2.90 | 9.0 | AB | AB | - | - | Stockbridge (63) |
| 7KKR[d] | | R25K | 3.11 | 6.2 | AB | - | - | - | Stockbridge (63) |

[a] R29K/E94C in *B. pertussis* and R25K in *E. coli* were introduced to enhance expression, selenomethionines (MSEs) were introduced to enhance phasing power (6). These replacements are functionally neutral. [b] pH where crystal grew. [c] Index of the chain that binds $F^-$. [d] $Br^-$ found at $S_{int}$.

#### 3.2 Other figures

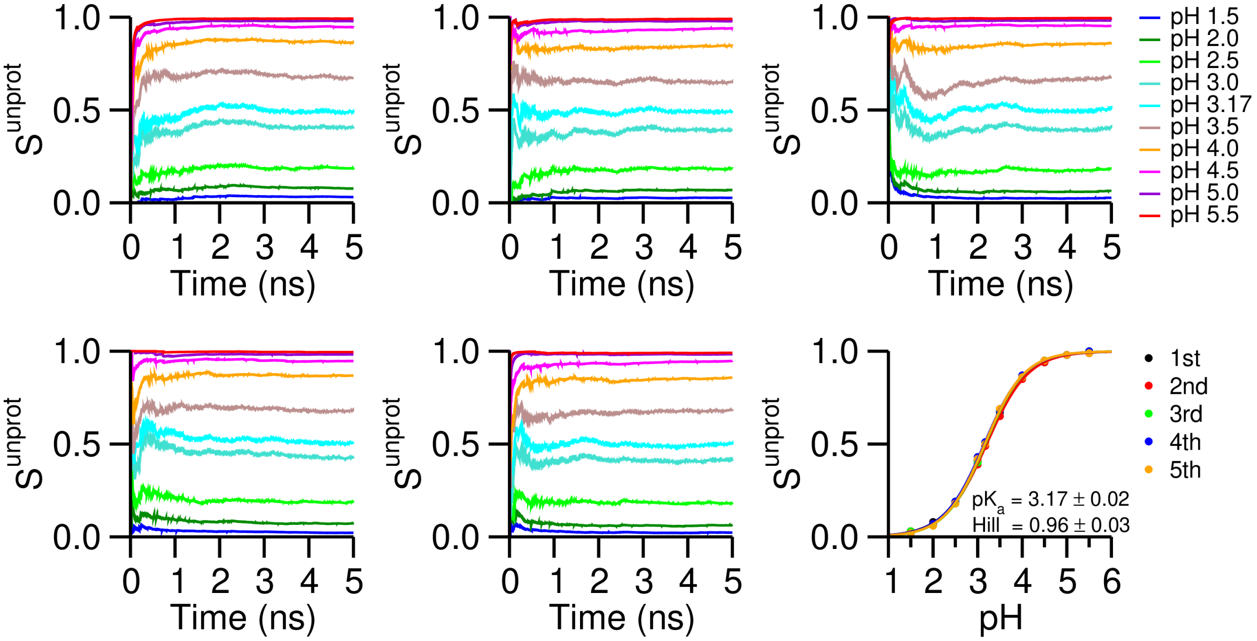


**Figure S10. Validation of the hybrid-solvent CpHMD parameters for HF.** 5 independent pH-REX CpHMD simulations were conducted to calculate the p$K_a$ of HF in aqueous. The titration curves are plotted in the bottom right panel with p$K_a$ and Hill coefficient $n$ reported (mean ± standard deviation). Other panels plot the convergence of the unprotonated fraction $S^{unprot}$ cumulatively calculated *vs*. time.

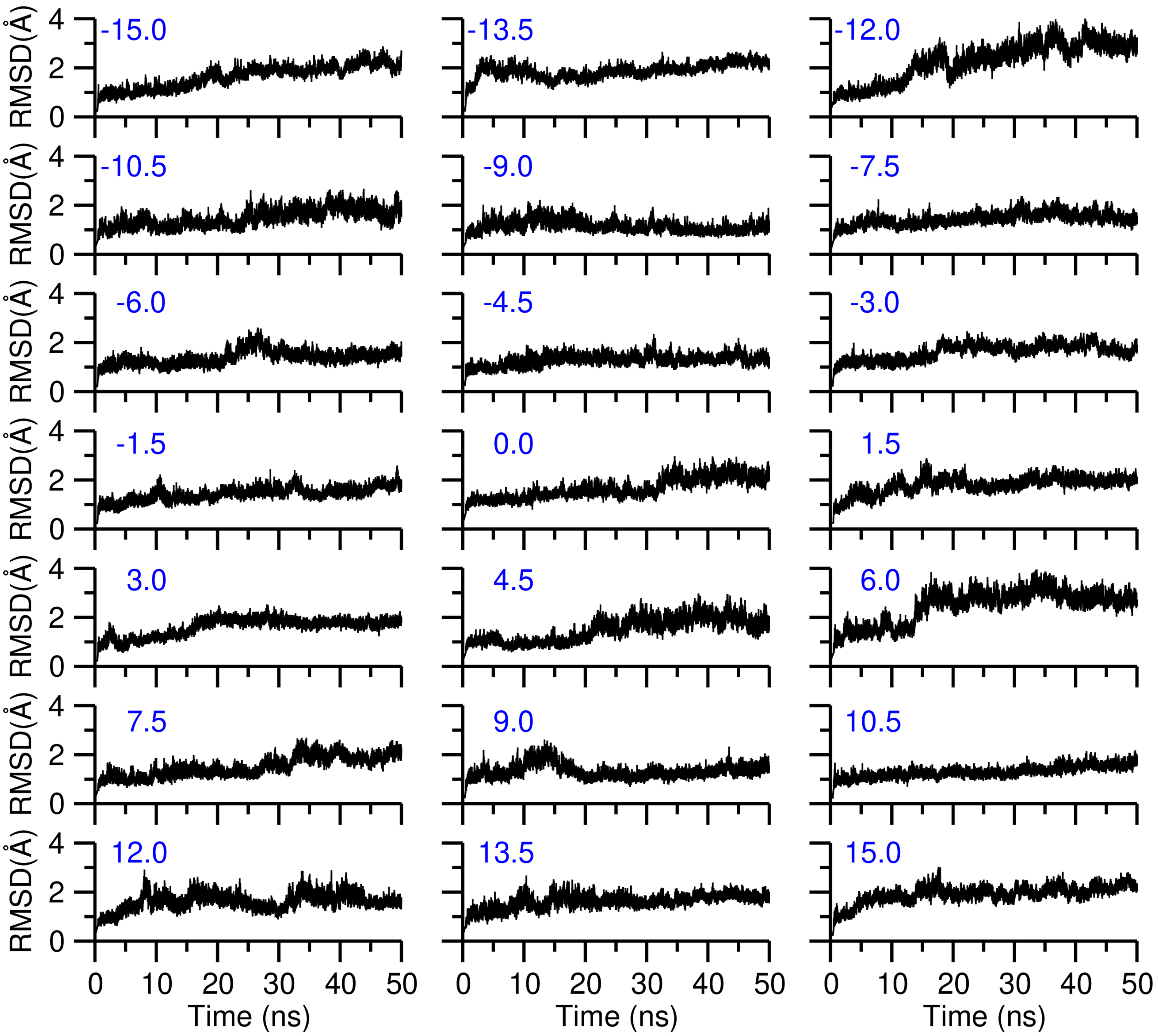


**Figure S11. System relaxation in the CpHMD equilibration.** Time series of the backbone RMSD (root mean square deviation) of Fluc-Ec2 are plotted for 21 replicates with the $F^{-}$ in monomer A restrained at various locations along the membrane normal (blue numbers at the top left corner of each panel, unit: Å).

### Appendix I: CHARMM topology and parameter for fluoride ion

```
* Toppar stream file for fluoride ion
* Parameters reported in Senn et al., J. Am. Chem. Soc., 2005, 127(39), 13643-13655
* Prepared by Zhi (Shane) Yue, Gregory A. Voth lab, UChicago, Dec 2019
*

!test "append" to determine if previous toppar files have been read and
!add append to "read rtf card" if true
set nat ?NATC
set app append
!We're exploiting what is arguably a bug in the parser. On the left hand side,
!the quotes have priority, so NAT is correctly substituted. On the right hand
!side, the ? has priority and NATC" (sic) is not a valid substitution...
if "@NAT" eq "?NATC" stop

read rtf card @app
* Topology for fluoride ion
*
36  1

MASS   501 FLA   18.99800 ! Fluoride ion

RESI FLA         -1.000 ! Fluoride ion
GROUP
ATOM FLA  FLA    -1.000
PATCH FIRST NONE LAST NONE

END

read para card flex @app
* Parameters for fluoride ion
*

ATOMS
MASS   501 FLA   18.99800 ! Fluoride ion

BONDS

ANGLES

DIHEDRALS

IMPROPER

NONBONDED

FLA    0.000000  -0.090000     1.810000 ! Fluoride ion
                                        ! from B. Roux via P. Jordan, dG = -111.8 kcal/mol

END
RETURN
```

### Appendix II: CHARMM topology and parameter for hydrofluoric acid

```
* Toppar stream file for hydrofluoric acid
* Parameters reported in Laage et al., Chem. Phys. Lett., 405 (4-6), 453-458
* Prepared by Zhi (Shane) Yue, Gregory A. Voth lab, UChicago, Jan 2020
*

read rtf card append
* Topology for hydrofluoric acid
*
36 1

MASS   502 FGA4  18.99800 ! F in hydrofluoric acid
MASS   503 HGA8   1.00800 ! H in hydrofluoric acid

RESI HF           0.000 ! Hydrofluoric acid
GROUP
ATOM F1   FGA4   -0.567 ! copied from Laage et al.
ATOM H1   HGA8    0.567

BOND F1 H1

DONOR H1 F1
ACCEPTOR F1

PATCH FIRST NONE LAST NONE

END

read param card flex append
* Parameters for hydrofluoric acid
*

ATOMS
MASS   502 FGA4  18.99800 ! F in hydrofluoric acid
MASS   503 HGA8   1.00800 ! H in hydrofluoric acid

BONDS
FGA4 HGA8  1000.000    0.9260 ! Kb set as a large arbitrary number, rigid H-F bond used by Drs. Laage & Hynes
                              ! b0 copied from Laage et al.

ANGLES

DIHEDRALS

IMPROPER

NONBONDED

FGA4   0.000000  -0.061000     1.559000 ! F in hydrofluoric acid, copied from Laage et al.
HGA8   0.000000   0.000000     1.320000 ! H in hydrofluoric acid, copied from HGA6, but epsilon made zero
                                        !                         personal communication with Dr. Laage

END
RETURN
```

### Appendix III: CHARMM hybrid-solvent CpHMD parameters for hydrofluoric acid

```
* States file for PHMD
* Charges reported in Laage et al., Chem. Phys. Lett., 2005, 405(4-6), 453-458
* Parameterized by Zhi (Shane) Yue, Gregory A. Voth lab, UChicago, January 2020
* Syntax:
* RESNAME MODEL_PKA PARA PARB
*  ATOM_TYPE CHARGE(1) CHARGE(2) [RAD(1) RAD(2)]
*

! -------------------------
! Experimental pKa: 3.165 +/- 0.007
! Measured at 298.15 K, 1 atm, 0.0001-0.01 M NaF using potentiometry
! Check Kresge et al., J. Phys. Chem., 1973, 77(6): 822-825
! -------------------------
! 5 independent pH-REX hybrid-solvent CpHMD runs
!
! CHARMM version: c42b2
! C22+CMAP CHARMM Force Field
! Truncated octahedron box, 32 Angstrom
! 1 HF, 801 CHARMM-modified TIP3P waters, no ions
! 298.00 K (Nosé-Hoover), 1 atm (Langevin piston)
!
! GB settings:
! gbsw hybrid sgamma 0.000 nang 50 conc 0.000 temp 298.00 -
!      sele SOLUTE end
!
! Nonbonded settings:
! nbond elec atom cdie vdw vatom vswitch -
!       ctonnb 10.0 ctofnb 12.0 cutnb 14.0 cutim 16.0 -
!       ewald pmew fftx 32 ffty 32 fftz 32 kappa 0.34 spline order 6 -
!       inbfrq -1 imgfrq -1
! -------------------------
! pKa(calc) = 3.17 +/- 0.02 (avg. +/- std. over 5 runs)
! n  (calc) = 0.96 +/- 0.03 (avg. +/- std. over 5 runs)
!
! Fraction of mixed state: 0.12 +/- 0.03 (avg. +/- std. over all pH's from 5 runs)
! -------------------------

HF  3.17 -90.5946 0.0986996 2.0
F1  -0.567 -1.000         !  H--F   <--->   F-
H1   0.567  0.000 1.0 0.0 !

END
```
